# Cross-study metagenomics analysis reveals distinct microbial signatures of urinary tract infections

**DOI:** 10.64898/2026.06.17.732905

**Authors:** David Giron-Villalobos, Constance Schultsz, Alan J. Wolfe, Marjon G.J. de Vos, Janneke H. H. M. van de Wijgert, Bas E. Dutilh, Michael F. Seidl

**Affiliations:** Theoretical Biology and Bioinformatics, Department of Biology, Utrecht University, Padualaan 8, 3584 CH, Utrecht, the Netherlands; Department of Global Health, Amsterdam Institute for Global Health and Development, Amsterdam UMC, University of Amsterdam, Amsterdam, the Netherlands; Department of Microbiology and Immunology, Stritch School of Medicine, Loyola University Chicago, 2160 S. 1st Ave, Maywood, IL 60153, USA; GELIFES, University of Groningen, Groningen, the Netherlands; Julius Center for Health Sciences and Primary Care, University Medical Center, Utrecht University, Utrecht, The Netherlands; Institute of Biodiversity, Faculty of Biological Sciences, Cluster of Excellence Balance of the Microverse, Friedrich Schiller University, Rosalind Franklin Strasse 1, 07743 Jena, Germany

**Keywords:** Urobiome, UTI, metagenomics, *E. coli*

## Abstract

**Background:** Urinary tract infections (UTIs) represent a major public health concern, increasingly complicated by rising antibiotic resistance, diminishing treatment efficacy, and increasing prevalence of recurrence. The urinary tract microbiome (urobiome) remains poorly characterized, despite its potential role in UTIs.

**Results:** To provide a comprehensive overview of the urobiome, we here integrated seven publicly available shotgun metagenomics studies, linking microbial composition to clinical infection status or diagnostics. Community-level analyses revealed distinct urobiome clusters, defined by one predominant bacterial taxon. Genome-resolved metagenomics allowed recovery of high-quality metagenome-assembled genomes (MAGs), enabling phylogenetic reconstruction, prediction of pathogenic potential, and profiling of antimicrobial resistance genes across multiple taxa. We then analyzed the pangenome of the clinically significant *Escherichia coli* species and found that its genomic variation is driven more by phylotype than isolation source or UTI status, supporting a model of opportunistic infection.

**Conclusions:** Taken together, our analyses represent a systematic, cross-study view of the urobiome that emphasizes the ecological complexity of the urobiome and the importance of integrating functional and phylogenetic information when studying UTIs.

## BACKGROUND

Microbiomes inhabit diverse niches within the human body [1]. They are increasingly recognized as a major factor impacting human health and disease [2]. Variations in the microbiome, specifically in the composition, abundance, and diversity of microbes in the community, have been implicated in the pathogenesis of acute and chronic diseases [3]. For example, alterations in the gut have been associated with cardiometabolic disorders, hypertension, neuropsychiatric disorders, cancer, or autoimmune diseases [4]. Similarly, skin and vaginal microbiomes have been extensively investigated in the context of disease [5,6]. Thus far, however, less attention has been paid to the role of the human microbiome in infectious diseases that have traditionally been attributed to a single pathogen [7]. Conversely, there is growing evidence that microbial communities can also positively influence disease outcomes [1]; for example, by directly improving immune responses or by modulating the interactions between the community and invading pathogens.

Until recently, the urinary tract was considered sterile in the absence of infection [8]. This view has changed with the establishment of enhanced culture methods and sensitive high-throughput sequencing technologies [9]. The urinary tract, like other body sites, has a resident microbiome, also referred to as the urobiome [10], which differs between males and females, and across age groups [11,12]. The adult male urobiome is often characterized by a high relative abundance of *Streptococcus,* other lactic acid bacteria, and/or *Corynebacterium* [13,14]. In contrast, the urobiome of post-pubertal pre-menopausal females is generally described as overlapping with the vaginal microbiome with high abundance of *Lactobacillus* species and lower abundances of other bacteria that are also often detected in the vagina. Moreover, peri-and post-menopausal women differ in their urobiome composition [15,16], likely caused by changes in the estrogen production associated with those life stages [17]. Due to its anatomical position, the urobiome has been linked to both the vaginal and gut microbiome, which are often considered potential reservoirs for uropathobionts [18], and recent studies have started to uncover possible mechanisms of microbial transfer between these body sites [19]. To further investigate the role of urobiome composition in human health and disease, genomics researchers have proposed the concept of urotypes using computational clustering of the bacterial taxa present in urine samples [20]. For example, urotypes predominated by *Lactobacillus*, *Gardnerella, Escherichia*, *Prevotella*, *Streptococcus*, or *Staphylococcus* have been described across different individuals [16,20]. While this framework provides a useful starting point to characterize urobiome composition across individuals, its clinical relevance remains uncertain [13]. In particular, it is currently unknown how specific community state types or urotypes are associated with urinary tract infections (UTIs) and its diagnosis, as current diagnostic guidelines rely on patient-reported symptoms and/or clinical microbiology culture results [21].

UTIs are among the most common bacterial infections affecting humans, either in the urethra, bladder, prostate, or kidneys [22]. It predominantly affects adult females, and approximately one in two females will experience at least one UTI in their lifetime, compared to only one in ten men [23]. Recurrent urinary tract infections (RUTIs) are also more prevalent in females, where 20-30% experiencing a recent UTI will also experience recurrence within six months, and this risk rises to 50% among postmenopausal females [24,25]. UTIs are common in otherwise healthy individuals, but are even more common in individuals with risk factors, such as pregnancy, metabolic disease, or immunosuppression [26].

*Escherichia coli* is the most common uropathobiont, accounting for over 75% of infrequent UTIs in healthy women using current non-molecular diagnostic methods [22]. However, its prevalence decreases after menopause (∼50%), reflecting a shift to other uropathobionts [27,28]. More recently, and thanks to more sensitive methods of detection and identification, other bacteria have emerged as potential uropathobionts, especially members of the *Aerococcus urinae* species complex, *Actinotignum schaalii,* and *Streptococcus anginosus* [29]. While some of these potential uropathobionts have well-characterized virulence mechanisms such as adhesins, biofilm formation, and iron acquisition [30], we know surprisingly little about their interactions with other members of the urobiome, including during UTI. This is especially relevant given that most bacteria taxa implicated in UTIs also are frequently detected in urine samples from individuals without UTI symptoms [13,31].

Experimental evidence suggests that urobiome composition, in addition to uropathobiont abundance, may explain susceptibility, progression, and treatment outcomes of UTIs [32]. In most clinical settings, uropathobiont identification still relies on culture-based methods that have been optimized for *E. coli* [33,34]. These methods, however, often fail to capture bacteria that require other growth conditions than *E. coli*, including slow growing, fastidious, and anerobic bacteria [35]. Microbiome sequencing represents an essential approach to overcome these limitations. To date, urobiome studies have primarily used 16S gene rRNA amplicon sequencing, which offers high sensitivity in determining confidence bacterial taxonomy, generally at the genus level [36]. Shotgun metagenomic sequencing also enables high-resolution taxonomy but has the additional benefit of providing insight into the encoded functions [10]. To our knowledge, no meta-analysis of metagenomic urobiome studies, including prediction of functions using microbial composition and associations with UTIs, has yet been conducted. Here, we performed such meta-analysis using publicly available metagenomic datasets. We then performed in-depth lineage-specific functional analyses on *E. coli*, the most commonly detected bacterial species associated with UTI diagnoses.

## MATERIAL AND METHODS

### Source data and quality control

To characterize the composition of the urobiome, we collected 433 publicly available whole-genome metagenomic sequence samples from NCBI (BioProjects IDs: PRJNA700071, PRJNA679884, PRJNA385350, PRJNA801448, PRJNA400628 and PRJNA680735) and ENA (Project accession: PRJEB45363). Raw reads were processed with the ATLAS metagenome pipeline v.2.18.1 [37] for quality control, assembly, and binning. Specifically, ATLAS employs BBTools [38] for quality filtering, metaSPAdes [39] for assembly into contigs, and vamb [40] for MAG binning. We assessed the quality of the MAGs using CheckM2 [41].

### Taxonomic composition of the urobiome

To determine the taxonomic composition of the urobiome, we applied the Metaphlan v4.0 [42] pipeline to classify reads against a clade-specific marker genes database based on the NCBI genomes database. We used a Snakemake [43] workflow to process multiple samples in parallel. Taxonomic profiles were aggregated and filtered to remove low-abundance taxa (<0.03 relative abundance). Subsequently, we used the phyloseq v1.52.0 package [44] in R to organize data at the genus level and integrate across studies. Next, we applied the mia v. 1.16.0 package [45] for relative abundance calculation; relative abundance values were transformed using CLR transformation. We then used vegan v2.7-1 package [46] to assess alpha and beta diversity with principal components analysis (PCA). Data was visualized with ggplot2 v3.5.2 [47].

### Phylogenetic tree of the urobiome

To explore evolutionary relationships in the urobiome, we reconstructed a bacterial phylogeny from the MAGs using the GTDB-Tk v2.4.0 [48] de *novo* workflow with default parameters. For the taxonomic annotation, we used a curated taxonomy classification version using BAT [49] against GTDB r220 [50] and kMetaShot [51] against RefSeq genomes NCBI. To estimate the number of reads mapped back to the MAGs, we followed the CAT/BAT/RAT pipeline [49,52]. Phylogenies were inferred from 120 bacterial marker genes, with Bacteroidota designated as the outgroup. Visualization and annotation of the tree were performed using ggtree v3.16.2 [53] and ggtreeExtra v1.18.0 [54] in R.

### Determination of pathogenicity capacity and antimicrobial resistance genes

The pathogenicity capacity of MAGs was assessed with PathogenFinder2 v0.5.0 [55], which predicts the likelihood of a bacteria being pathogenic in humans based on their genome content. AMR presence in genomes was performed using RGI v6.0.5 querying the database CARD v3.2.7 [56].

### *E. coli* phylogenetic analysis

Given the clinical relevance of *E. coli* in the urinary tract, we conducted a detailed comparative analysis. We downloaded 54,000 publicly available *E. coli* genomes from the BV-BRC database (June 2025) (**Tab. S5**) [57] and combined them with the *E. coli* MAGs identified in our study, as designated by BAT and kMetaShot. To determine their phylogroups, we applied ClermonTyping v24.02 [58] as phylogroups are often associated with distinct ecological niches and pathogenic potential [58]. To place our MAGs within the global *E. coli* diversity, we performed a clustering analysis using PopPUNK v2.7.6 [59], which grouped genomes into 1,916 distinct clusters. A representative from each cluster was selected and a phylogenetic tree was constructed with Mashtree v1.4.6 [60], including both reference genomes and our MAGs. Tree visualization was performed with ggtree v3.16.2 [53] and ggtreeExtra v1.18.0 [54].

### Pangenome construction, functional annotation, and virulence factor profiling

To investigate genomic variation among *E. coli* genomes, we constructed a pangenome using Panaroo v1.5.2 [61] with default parameters. Genes present in ≥95% of genomes were defined as core, those present in 15–95% as shell, and those in <15% as cloud genes. Based on the resulting presence-absence matrix, we performed a genome-wide association study (pan-GWAS) using Scoary v1.6.16 [62] to identify genes significantly associated with strain-level traits (e.g., phylotype or isolation source). Genes were considered significant if they met the following criteria: Bonferroni-adjusted p < 0.05 and both specificity and sensitivity > 75%. To estimate the openness of the pangenome, we calculated the α parameter from Heaps’ law using the R packages micropan v. 2.1 [63] and vegan v. 2.7-1 [46], and then visualized the results with ggplot2. Functional annotation of all pangenome genes was performed with eggNOG-mapper v2.1.13 [64], database eggNOG v5.0.2 [65], and functional categories were assigned based on COG annotations. Statistically significant genes identified by Scoary were mapped to these functional categories to assess their biological roles. To quantify the distribution of virulence-associated genes, we used the VFDB (downloaded November 2025) [66]. Using diamond v. 2.0.15 [67], a database was built from VFDB protein sequences and predicted protein sequences from the pangenome were queried against this database with blastx using the “fast” mode to identify homologous virulence genes. Detected genes were assigned to 14 categories of virulence factors, defined in VFDB. For genomes isolated from human sources (gut, respiratory, blood, urine, or other), we calculated the proportion of virulence genes per category relative to total gene count for each genome. To evaluate differences among isolation sources, we performed a Kruskal-Wallis’s test followed by post hoc Dunn’s test for pairwise comparisons. Proportions were visualized using a beeswarm dot plot implemented with the ggbeeswarm package, with samples ordered by the smiley factor.

## RESULTS AND DISCUSSION

### Exploring the urobiome with publicly available shotgun metagenomics datasets

To systematically describe the composition of the urobiome in the presence or absence of a UTI diagnosis, we collected and reanalyzed publicly available shotgun metagenomic sequencing datasets from seven independent studies [15,19,68–72]. Six of these studies were conducted in the United States, and one in the Netherlands. These studies differed in sampling methodology (e.g., midstream urine or catheter collection), as well as in protocols for sample handling, storage, and DNA extraction. All sequencing datasets were generated with different short-read sequencing platforms (**Tab. S1**). Samples were typically collected either as part of UTI diagnostics or of urobiome-focused studies. Based on clinical classifications provided in the original publications, we divided participants into four UTI status groups (**Fig. 1A**, **Tab. S2**). Patients with a culture-confirmed UTI on growth media that selects for common uropathobionts (growth > 10^4^ CFU/mL) [73,74] were classified as ‘culture-confirmed UTI’ (ccUTI). Patients that were recruited as controls (i.e., did not have any symptoms at the time of sampling, no RUTI diagnosis, and no chronic urinary tract disease history) were classified as ‘negative UTI’ (negUTI). We also defined two additional groups that contained a mixture of patients. Patients with a probable/possible UTI based on reported symptoms but no, little, or atypical bacterial growth on culture media, as well as those diagnosed with ‘recurrent UTI’ (RUTI) were classified as status ‘unclear related to UTI’ (uncUTI). Finally, patients with chronic urinary tract disease history, such as interstitial cystitis or overactive bladder, but with no or unknown symptom-status at the time of sampling were classified as status ‘unclear chronic disease’ (uncCD). After excluding mock communities, 433 samples were retained for further analyses, from 421 different patients (**Fig. 1A**, **Tab. S2**). From the available metadata, we identified 289 samples as being derived from females and 58 from males, and for 86 no sex was reported.

**Figure 1.**
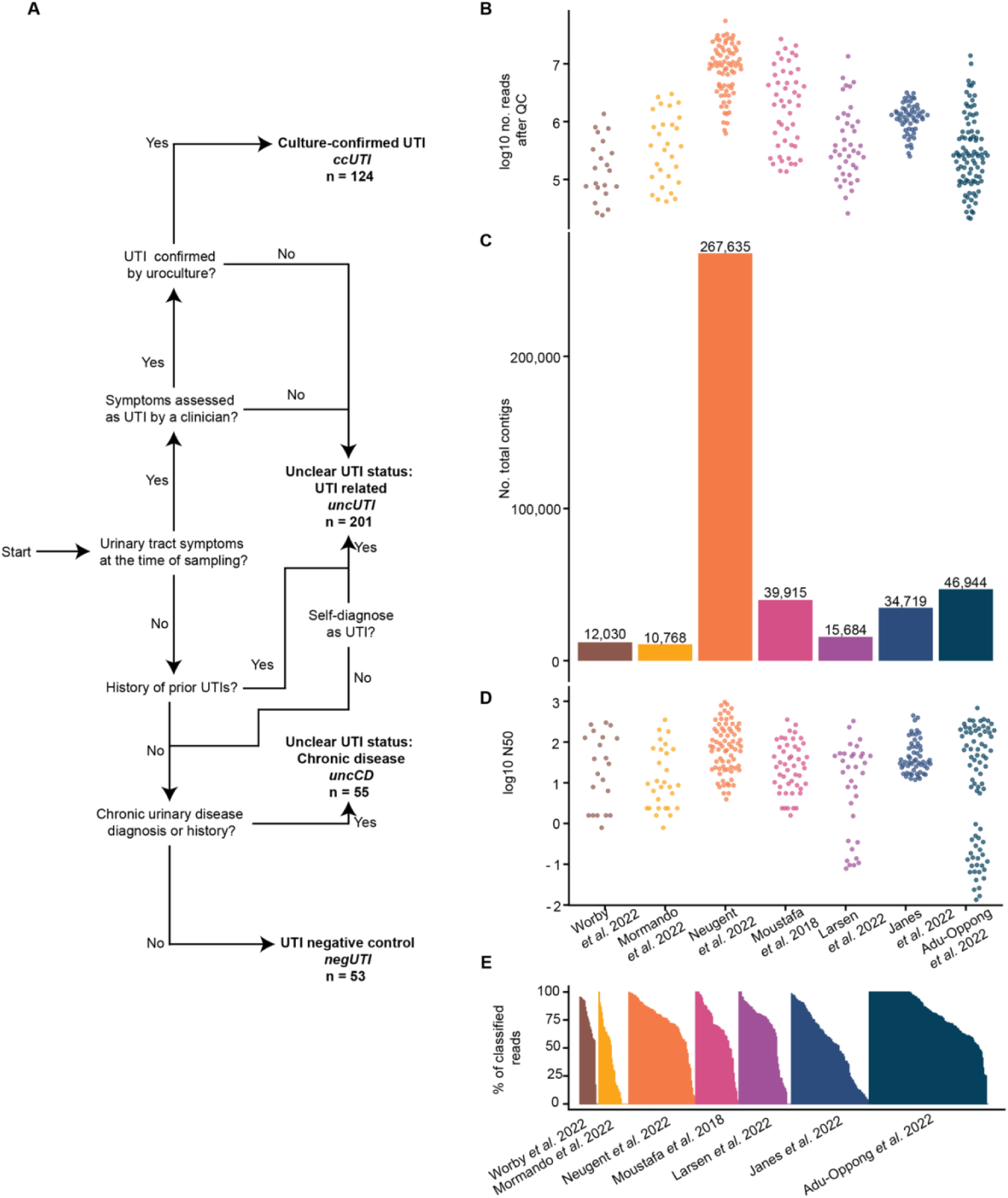
Quality control and assembly statistics across urobiome metagenomics studies reveal different sequencing results across 433 samples. **A)** Samples were grouped into four distinct groups ccUTI, negUTI, uncUTI, and uncCD, following this decision-making workflow using conditions found in the original publications (for details, see **Tab. S2**). **B)** The distribution of the number of quality-controlled paired reads (log_10_ transformed) after processing shows clear differences in sequencing yield across studies. **C)** Contigs assembled at sample level with metaSPAdes (length cutoff 1,000 bp) were aggregated per study, highlighting variability in assembly outcomes. **D)** N50 values (log_10_ transformed) per sample are shown, aggregated by the study. Higher N50 values indicate more contiguous assemblies, with clear variation observed across studies. **E**) Fraction of reads classified at the species rank with Metaphlan per sample, colored and grouped by study. Samples with high classification rates were observed in all studies, despite differences in sequencing methods.

### Urobiome signals can be detected across studies despite technical heterogeneity

Due to the low microbial biomass typical of urine samples from individuals without a UTI, sequencing the urobiome to detect microbial signatures of the resident microbes is challenging [36]. We thus first sought to assess the capacity of shotgun metagenomic sequencing to identify and classify the microbial composition of the urobiome. To this end, we obtained the raw sequencing data from public repositories and ran the same analytic pipeline on all samples (see Methods). We performed quality control by removing duplicate and human-derived sequencing reads, eliminating on average around 1.16M +/- 1.61M read pairs per sample, yielding on average a total of 3.74M +/- 6.89M quality-filtered paired reads. The number of high-quality reads varied substantially between studies (**Fig. 1B**), reflecting differences in sampling, DNA extraction, library preparation, and variations in Illumina sequencing platforms (**Tab. S1**). Notably, samples from the study by Neugent and colleagues [15] contained consistently more quality filtered reads (∼10 million total paired reads per sample) than samples from the other studies (2∼10 million paired reads, **Fig. S1**).

To obtain microbial contigs from each individual sample, we used the ATLAS pipeline to process the quality-filtered reads [37]. We assembled the reads with metaSPAdes, yielding on average 1,178 +/-2,102 contigs per sample. As expected, the study by Neugent and colleagues [15] consistently produced more contigs with higher N50 values (**Fig. 1C, 1D**). Across samples, between 7.8 to 99.8% of reads mapped back to the assembled contigs with a mean of 82.3%, quantifying to what extent the contig data reflects the sequenced community.

### Taxonomic profiling reveals urobiome signatures across different cohorts

To determine the composition of the urobiome in the sampled individuals, we used Metaphlan [42] for taxonomic classification at the read level. Despite differences in extraction protocols and total read yield, a substantial proportion of reads were successfully classified across most samples (**Fig. 1E**), with an average classification rate at the species rank of 69.6% (ranging from 2.85% to 100%). We observed high classification rates in the studies conducted by Neugent and colleagues, Worby and colleagues, as well as Adu-Oppong and colleagues [15,72,75], with an average of 75% reads classified, while other studies showed more variation, averaging between 50 - 70% (**Fig. S2**). Notably, this variation was not associated with study-specific protocols, suggesting that other aspects besides technical differences determine the classification success. Importantly, the fact that a substantial fraction of reads could be classified in most studies suggests the presence of a detectable microbial signal within the urinary tract, despite urine being a low microbial biomass environment [36].

### Urobiome composition reveals community clusters

To provide a comprehensive microbial profile of the urobiome, we integrated the taxonomic annotations from the seven metagenomic studies using relative abundances. To minimize batch effects and methodological variability, we applied centered log-ratio (CLR) transformation, followed by hierarchical clustering of the 433 samples on genus-rank relative abundances (**Fig. 2A**). This cross-study analysis clustered the 433 samples into 13 distinct clusters by microbial community similarity. Twelve of these clusters were defined by one specific bacterial genus or species, while one was a mixture of taxa with no clear predominant member. We named the clusters based on the defining genus or species when specific taxon defined it, or as ‘mixed’. Five clusters contained high relative abundances of known uropathobionts, namely *E. coli, Klebsiella pneumoniae*, *Pseudomonas aeruginosa, Proteus mirabilis,* and *Enterococcus* [22]. Four clusters contained bacterial genera commonly associated with the vaginal, gut or skin microbiomes with varied species representatives (*Lactobacillus*, *Gardnerella*, *Bifidobacterium*, and *Cutibacterium* [26] and three clusters contained known pathobionts that can occur in multiple body niches (*Staphylococcus*, *Streptococcus*, or *Citrobacter*). Importantly, most clusters were found in three or more different studies (**Fig. 2A**), suggesting limited study-dependent biases. Some of the identified clusters have been previously described, for example, *Escherichia-Shigella, Gardnerella, Lactobacillus, Streptococcus, Staphylococcus,* and [20,76–79]. Other studies also identified additional clusters with high relative abundances of *Acinetobacter, Anaerococcus*, *Corynebacterium*, and *Sphingomonas.* Although we detected this data in our dataset, we did not observe corresponding clusters.

**Figure 2.**
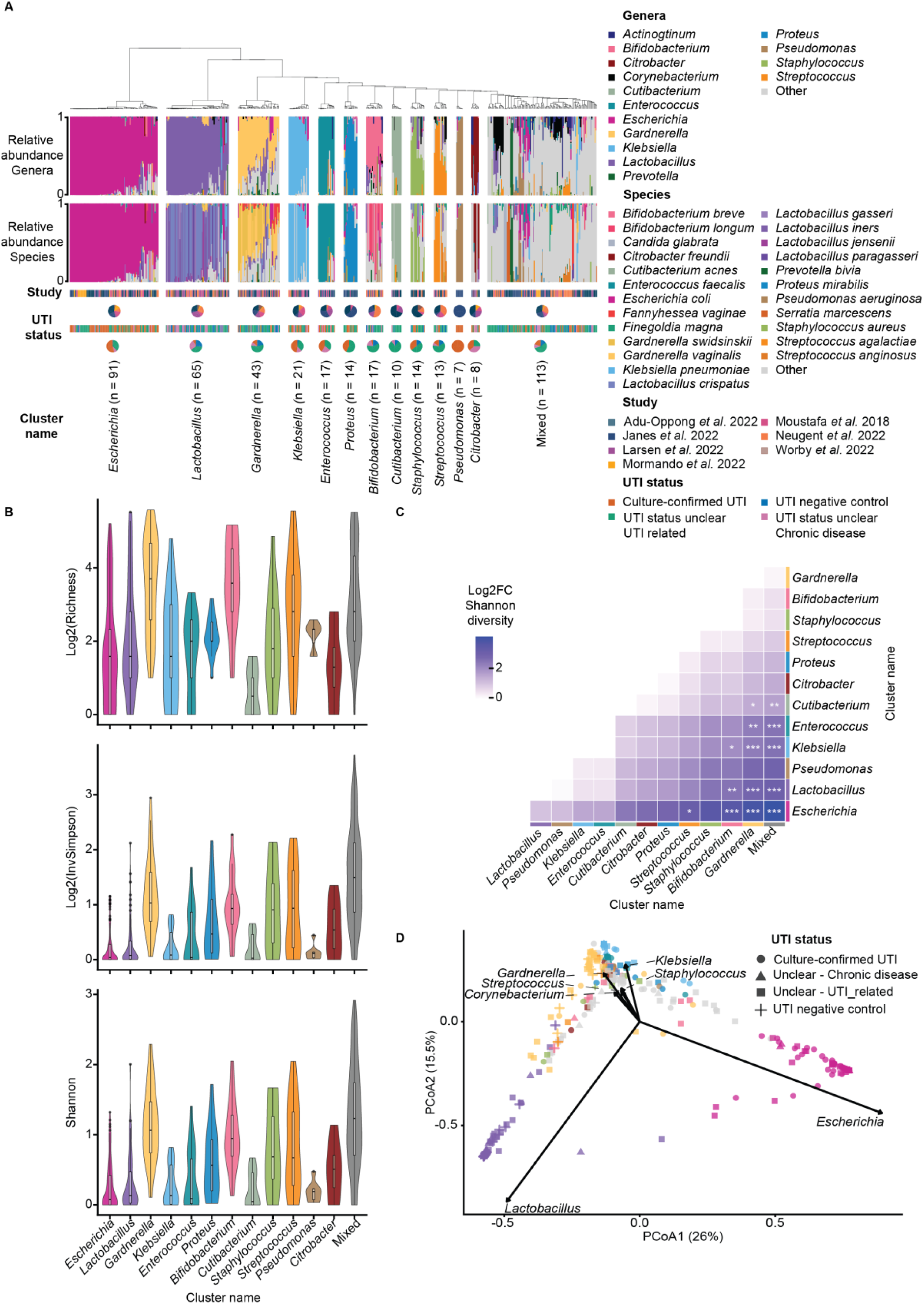
Urobiome composition varies across individuals with and without UTI. **A)** Genus- and species-rank relative abundances across all samples after centered log-ratio (CLR) transformation to normalize study variability. Samples were hierarchically clustered based on compositional similarity and grouped into 13 clusters. Associated metadata for each sample, including study and UTI status at the time of sampling, are shown below the abundance plot. Pie charts indicate the proportion of the different metadata categories represented in every cluster. Clusters do not align with UTI status or study. **B)** Alpha diversity (Richness, Inverse Simpson, and Shannon index) for the samples in each cluster identified in panel **A**. Violin and box plots show the distribution of diversity metrics and their median. Statistical significance among clusters was assessed (Kruskal-Wallis’s test, *p*< 0.05). **C)** Heatmap of pairwise log2 fold changes in median Shannon diversity between clusters. Statistical significance was evaluated using Dunn’s test with Bonferroni correction (\**p* < 0.05, \*\**p* < 0.01, \*\*\**p* < 0.001). **D)** Principal coordinate analysis (PCoA) based on Euclidean distances of the transformed relative abundances. Data points were colored by cluster, as shown in panel **A**. Each point represents a sample and is shaped by UTI status at the time of sampling. Samples did not cluster by UTI status or by study (**Fig. S3**), but by the relative abundance of specific genera (*Escherichia* and *Lactobacillus*).

To further characterize the bacterial composition of samples within and between these 13 clusters, we described their alpha- and beta-diversity. The *Escherichia*, *Klebsiella*, *Enterococcus*, *Cutibacterium*, and *Lactobacillus* clusters exhibited significantly lower alpha diversity than more heterogeneous clusters *Gardnerella*, *Bifidobacterium* and, mixed clusters (**Fig. 2B, 2C**). Principal Coordinates Analysis (PCoA) revealed that separation of samples was mainly driven by the relative abundance and prevalence of *Lactobacillus*, *Escherichia, Klebsiella,* and a mixture composed by *Gardnerella, Streptococcus, Staphylococcus,* and *Corynebacterium* (**Fig. 2D**), Moreover, no clear clustering was observed on age or by study (**Fig. S3**), suggesting that these factors have less impact on the composition of the urobiome. Notably, 54% of samples within the *Escherichia* cluster originated from individuals in the ccUTI group, whereas only 7% of samples in the *Lactobacillus* cluster corresponded to ccUTI (**Fig. 2A**). However, interpretation of these observations should be tempered with caution by considering differences in the diagnostic and sample selection methods used between UTI status groups. In the ccUTI group, microbial identification relied on growth under standard clinical microbiology culturing conditions, and these conditions are known to preferentially capture *Escherichia* and other Gram-negative uropathobionts. While the various ecological metrics that we presented provide valuable insights into community organization, further integration with clinical phenotypic, and experimental data is required to fully understand their biological and clinical relevance.

### Urobiome composition patterns associated with age and UTI status

To further investigate the relationship between urobiome composition, UTI status, and host phenotype, we focused on females because they were the most clinically well-characterized samples. This subset included ccUTI (n = 73), uncCD (n = 16), uncUTI (n = 83), and negUTI (n = 42) samples. We first assessed whether the female urobiome composition varied with age, as age represents a clinically relevant factor in female urinary tract health due to the decline in estrogen after menopause, which typically occurs around the age of 50 [16,80]. We evaluated the effects of age within the clinically best characterized ccUTI and negUTI groups by categorizing the urobiome composition within as predominance (> 50% of relative abundance) of the Gram-negative uropathobionts *Escherichia*, *Proteus*, and/or *Klebsiella*, predominance (> 50%) of *Lactobacillus* species, or “other” (all left-over samples) (**Fig. 3A**). Within the ccUTI group, the median age of the women with uropathobiont predominance (68 years, range: 21 - 99) did not differ significantly from the median age of the females in the other group (61 years, range: 19 - 87) (**Fig. 3B)**. However, the proportion of postmenopausal women with uropathobiont predominance (66.67%) was higher compared to the presence of another urobiome (33.33%). Within the negUTI group, no significant differences were found between the median age of the *Lactobacillus* predominance (60 years, range: 24 - 82) and another urobiome groups (60 years, range: 19 - 81) (**Fig. 3B**), while the proportion of postmenopausal women with *Lactobacillus* predominance group (31%) was lower comparing to females with other urobiome (69%). We next determined changes in alpha diversity across age and UTI status groups using linear regression analyses (**Fig. S4**). Overall, these analyses suggest an increase in richness with age within the ccUTI, uncUTI, and negUTI groups. In the negUTI group, Shannon diversity indices and age were positively associated (**Fig. S4**). To further explore age-related trends, we performed differential abundance analysis using MaAsLin3 [81], which fits multivariable linear models to identify associations between microbial features and continuous or categorical data; age and UTI status were included as covariates in the model. *Finegoldia magna* was positively associated with age, while *E*. *faecalis*, *Gardnerella vaginalis*, and *K*. *pneumonia* were negatively associated with age across all UTI status groups (**Fig. S5A**). Prevalence-based analyses, however, revealed a more complex picture; these analyses only considered the presence or absence of taxa, regardless of their relative abundance. Log-odds ratio estimates indicated that several taxa, including *Escherichia coli* and *E. faecalis*, were more likely to be present in women of older age, alongside the genera *Corynebacterium*, *Peptoniphilus*, *Anaerococcus*, *Actinotignum*, *Winkia*, *Finegoldia*, *Streptococcus,* and *Staphylococcus* (**Fig. S5B**). In contrast, taxa such as *Lactobacillus iners*, *Fanyhessea vaginae*, and members of the genus *Gardnerella* were more prevalent in women of younger age. Finally, differential abundance analysis identified *E. coli*, *E. faecalis*, and *K. pneumoniae* as significantly enriched in ccUTI samples compared to negUTI samples (**Fig. 3D**). Conversely, prevalence analyses suggested a higher occurrence of *Anaerococcus*, *Corynebacterium*, *Cutibacterium*, *Lactobacillus*, *Peptoniphilus*, *Staphylococcus*, *Varibaculum*, and *Winkia* in the control group. Together, these results provide a detailed view of how urobiome composition varies with both age and UTI status. We consistently observed an increase in richness with age in both negUTI and ccUTI groups, which may reflect a transition toward more complex urobiome communities in older individuals, irrespective of their clinical status. This is further supported by differential abundance and prevalence analyses, which indicate that the contribution and occurrence of multiple taxa vary across age. UTI status is also characterized by evident compositional differences: ccUTI samples were enriched in classical uropathobionts, whereas negUTI samples display a predominance of either *Lactobacillus* or exhibited more diverse microbial profiles. However, these patterns do not fully explain cases where no single uropathobiont predominates or where non-classical taxa arepresent. In particular, the role of mixed microbial communities and the potential contribution of taxa present in control-associated profiles but missing in ccUTI remain unclear. These observations highlight the need for further investigation into how shifts in community composition, beyond single-uropathobiont predominance, contribute to UTI development.

**Figure 3.**
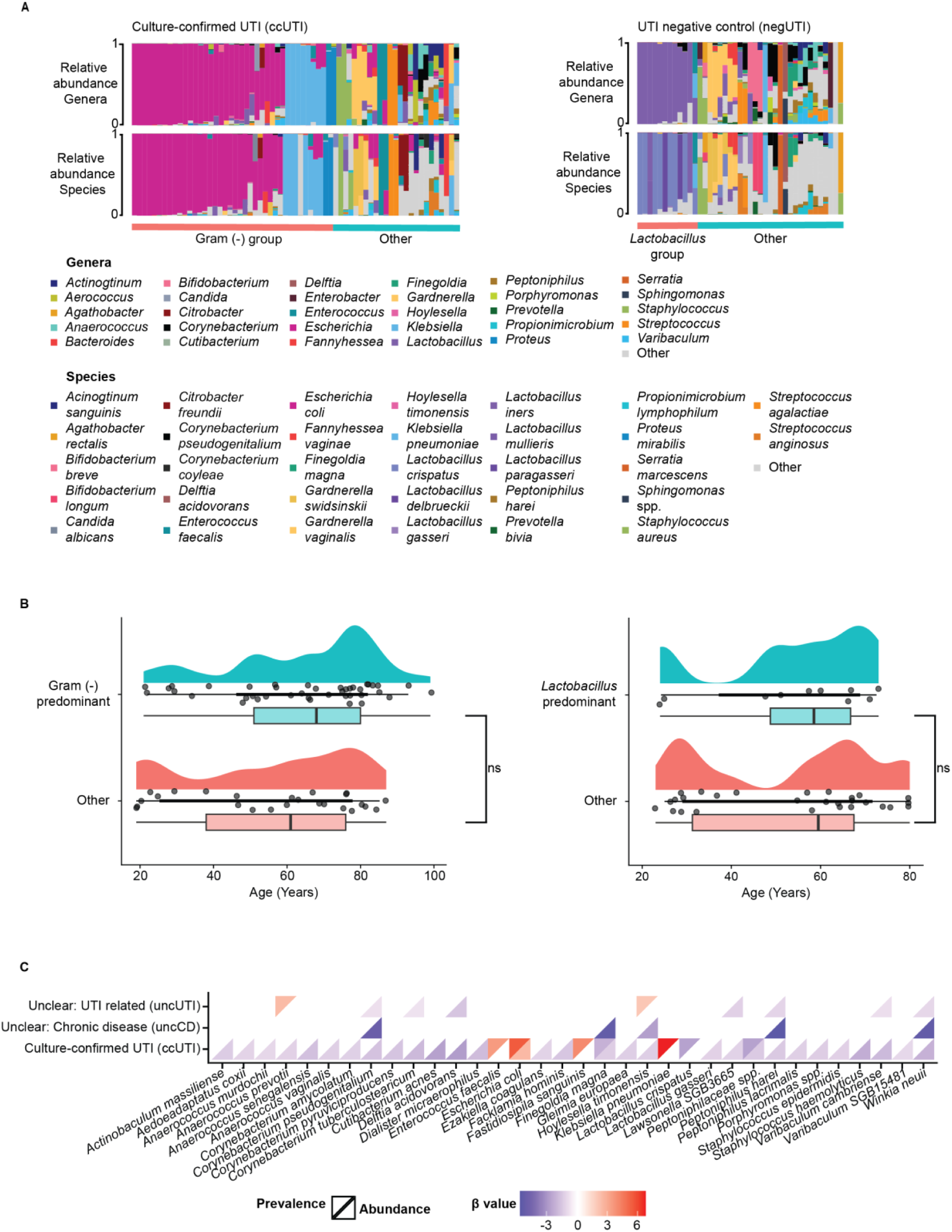
Urobiome composition patterns associated with age and UTI status. **A**) Genera- and species-rank bar plots display the relative abundances of samples from the ccUTI and negUTI groups. Samples are sorted according to their relative abundance of Gram-negative uropathobionts (ccUTI, left) and *Lactobacillus* species (negUTI, right), respectively, with all remaining samples labeled as “Other”. This highlights the importance of uropathobionts for ccUTI and *Lactobacillus* species for negUTI. Other, often mixed samples occurred in both groups. **B**) Distribution of age in the different sample groups shown as density plots (top) and boxplots (bottom). Boxplots display the median (center line), interquartile range (box) and data spread (whiskers), with individual points representing samples. Density plots illustrate the overall distribution of ages within each group. Median age is comparable between the groups of interest and “Other” (Wilcoxon test p>0.05). **C**) Differential abundance analysis between UTI status groups, using negUTI as a reference group. Negative coefficients indicate enrichment in the negUTI group, whereas positive coefficients indicate enrichment in the ccUTI group. Each square is divided into two triangles: the upper triangle represents differential prevalence values, and the lower triangle represents differential abundance values.

### Binning recovers high-quality metagenome-assembled genomes from urine

Shotgun metagenomics provides insights into the genomic content and functional potential of the urobiome. We therefore used the metagenomic data to reconstruct metagenome-assembled genomes (MAGs) from assembled contigs through sample-level binning, recovering a total of 1,072 sample-specific MAGs from all samples and datasets (size range: 4,194 – 11,401,458 bp, mean: 1,643,933 bp) (**Tab. S3**). We then assessed the MAG quality using standard thresholds for completeness and contamination [82], yielding 420 high-quality, 234 medium-quality, and 418 low-quality MAGs (**Fig. 4A, 4B**). While the number and quality of MAGs varied across studies, reflecting differences in sequencing depth and sample complexity, high-quality MAGs were nonetheless recovered from all datasets. This highlights that even studies with fewer or lower-quality MAGs can still provide valuable genomic information, although studies with higher sequencing volume did generate more high-quality MAGs. High-quality MAGs spanned a wide range of sizes (667,127 – 9,564,481 bp), emphasizing that genome-resolved metagenomics can capture diverse microbial genomes from urine.

**Figure 4.**
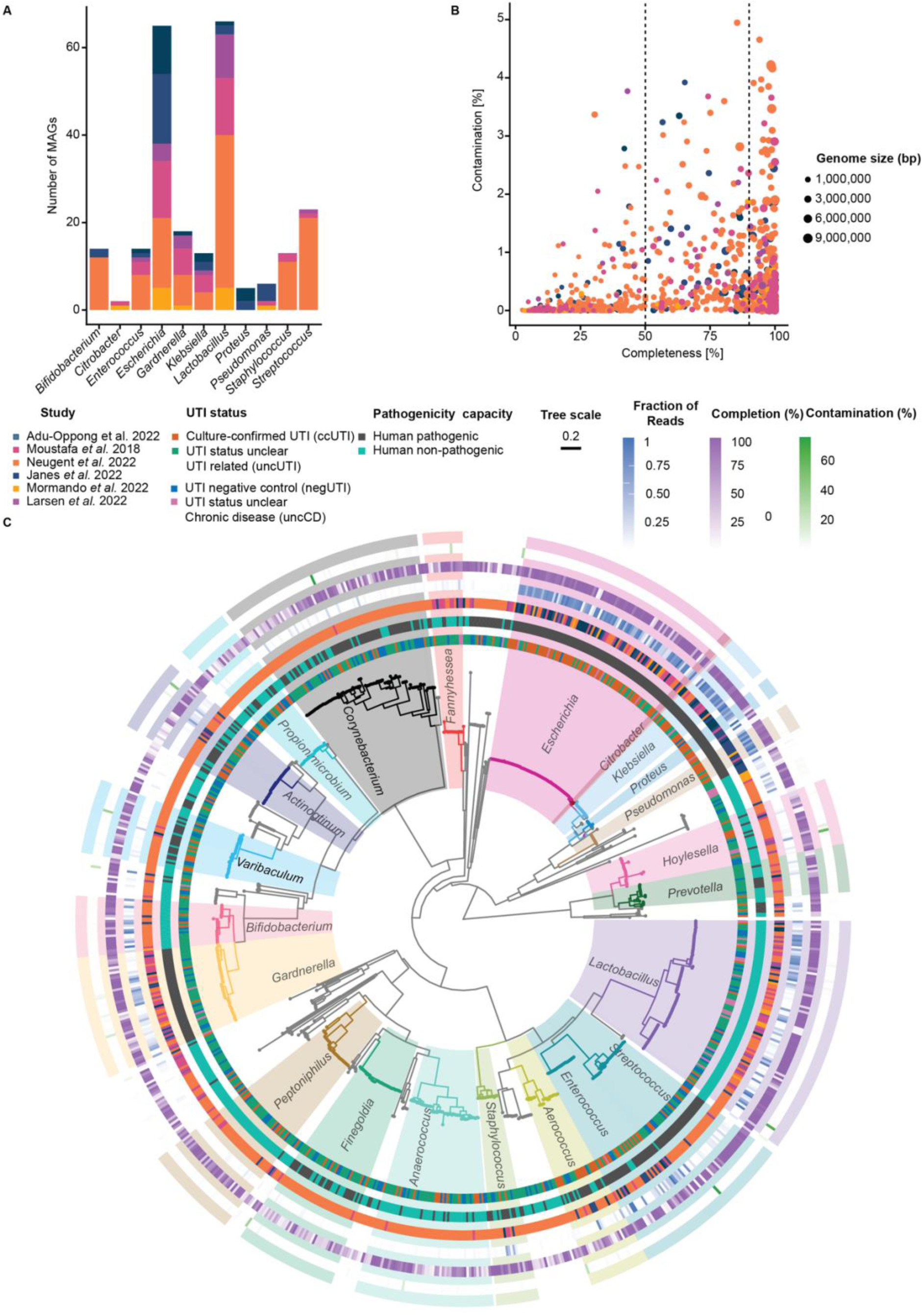
Phylogenetic analysis of the urobiome-derived metagenome-assembled genomes (MAGs) reveals diverse taxonomic groups. **A)** Top genera are consistently represented across studies despite variation in MAG recovery. Bars show the number of genomes, colored by the number of high-quality MAGs recovered for every study distributed by identified top genera. **B)** Distribution of MAG quality metrics and size, showing that there are high-quality MAGs in all samples. **C)** Phylogenetic tree of MAGs, colored by bacterial genus. Concentric rings display associated metadata, from inner to outer ring: UTI status, predicted pathogenicity, study of origin, and MAG quality: heat strips showing estimated completion, and contamination. Together, these features revealed a complex relationship between bacterial composition, predicted pathogenicity, and host phenotype.

### Metagenome-assembled genomes include heterogeneous microbial lineages with a diverse functional potential

To taxonomically classify MAGs, we then applied two complementary approaches. While BAT [49] offers a robust framework using the GTDB (v. r220) taxonomy, its coverage of clinically relevant urobiome taxa is currently limited (e.g. *Gardnerella* and related vaginal-associated taxa). To address this limitation, we also classified MAGs with kMetaShot [51] using the NCBI/RefSeq bacterial and archaeal database. BAT classified 96% MAGs at the genus rank and 50% at the species rank, respectively, whereas kMetaShot classified 88% at the genus rank and 65% at the species rank. While most MAGs originated from the study by Neugent and colleagues [15] (**Fig. 4A**), the most abundant genera (i.e. *Escherichia*, *Gardnerella*, and *Lactobacillus*) were recovered from multiple datasets (**Fig. 4A, Fig. S6, Tab. S3**).

To explore the phylogenetic diversity of the urobiome MAGs, we next reconstructed a bacterial phylogeny with GTDB-Tk [48], retaining 898 MAGs with at least 50% alignment coverage of the conserved protein markers used by GTDB-Tk for phylogenetic inference. We then used the combined taxonomic classification from BAT and kMetaShot to assign MAGs to taxonomic lineages. The resulting phylogenetic tree captures the heterogeneity of bacterial lineages present in the urobiome (**Fig. 4C**). In most bacterial genera, we observed several divergent clades, suggesting the presence of multiple species. As an exception to this rule, all *Escherichia* MAGs formed a single clade with very low diversity, suggesting that the number and diversity of urine-associated *Escherichia* species is much lower than that of other genera. This was further corroborated by comparing the phylogenetic distances of members in the same genus (**Fig. S7**). In addition to *Escherichia*, four others known uropathobionts (*Klebsiella*, *Proteus*, *Pseudomonas*, and *Enterococcus*) displayed a relatively low intra-genus divergence, the intra-genus distance was higher for bacteria commonly found in the vagina and the skin microbiome (e.g. *Gardnerella*, *Lactobacillus*, *Streptococcus*, and *Staphylococcus*) (**Fig. S7**). Together with our taxonomic profiling results (**Fig. 2**), these findings emphasize that the urobiome has a broad phylogenetic diversity beyond the known uropathobiont genera and species.

While certain genetic and phylogenetic patterns have been linked to uropathogenicity [83], they remain inconsistent and difficult to generalize [22]. To investigate the pathogenic capacity of our reconstructed MAGs, we applied PathogenFinder2 [84] to predict human pathogenic potential based on the presence of known virulence genes. MAGs predicted to be pathogenic aligned well with the classical uropathobionts *Enterococcus, Escherichia*, *Klebsiella*, *Proteus*, and *Pseudomonas* (**Fig. 4C**). *Escherichia* was predicted as pathogenic and was present in ∼ 40% of all samples, while other known uropathobionts showed a comparable pattern, even though their prevalence was considerably lower (**Fig. S8**). However, this analysis also revealed mismatches between predicted pathogenicity and UTI status. For several taxa, including *Actinotignum*, *Corynebacterium, Gardnerella, Finegoldia,* and *Streptococcus,* most MAGs were predicted to be pathogenic based on their genome content (**Fig. 4C, Fig. S8**), despite being recovered from negUTI patients. As expected, *Lactobacillus*, which is typically regarded as beneficial [85], was consistently predicted as non-pathogenic, with presence in ∼5% of the ccUTI group and in ∼30% of the negUTI group. These results suggest that bacterial pathogenicity genes are preferentially, but not exclusively, associated with specific taxonomic groups [86,87]. Our findings align with and extend the model of UTI pathogenesis by predominant uropathobionts with a vast diversity of both predicted pathogenic and non-pathogenic urobiome members [22].

### Antimicrobial resistance is widespread across urobiome communities

Interspecies interactions can shape microbial dynamics and antibiotic resistance [89,90], but are often overlooked when considering UTI treatment strategies [91]. To characterize antimicrobial resistance (AMR) within the urobiome, we screened for the presence of AMR-associated genes in all MAGs using RGI queried against the CARD database [56]. AMR genes were found in 35/82 bacterial genera (**Fig. 5, Tab. S4**). On average, *Pseudomonas* and *Escherichia* MAGs contained 55.4 +/- 1.9 and 43.6 +/- 11.2 AMR genes, respectively, whereas *Lactobacillus* and *Gardnerella* contained very few (1.5+/- 0.5 and 1.1 +/- 0.3 AMR gene per genome, respectively). The high number of detected AMR-associated genes likely reflects the nature of the annotation framework. For example, CARD includes determinants such as efflux pumps, whose presence may not confer clinically relevant resistance [92]. Additionally, it includes specific sequence mutations to *Escherichia*, *Klebsiella*, *Proteus*, and *Pseudomonas* that primarily reflect taxa-specific genetic resistance associations [93]. Recently, an analogous large comparative genomic survey of AMR found that *Escherichia* from diverse environmental and clinical sources carry on average 55 AMR genes [93]. Similarly, another study focused on bloodstream infection isolates, reported that *Pseudomonas* harbor 33, *Escherichia* 30, and *Lactobacillus* five AMR genes per genome [94], suggesting that uropathobionts are generally enriched for AMR-associated genes. We hypothesize that this enrichment partly reflects selection by antibiotic treatment of individual patients, partly by antibiotics in the greater anthroposphere, and the potential capacity of uropathobionts to acquire and maintain AMR genes through horizontal gene transfer. However, because our analyses did not distinguish between chromosomal and mobile AMR elements, the relative contribution of these mechanisms remains unclear.

**Figure 5.**
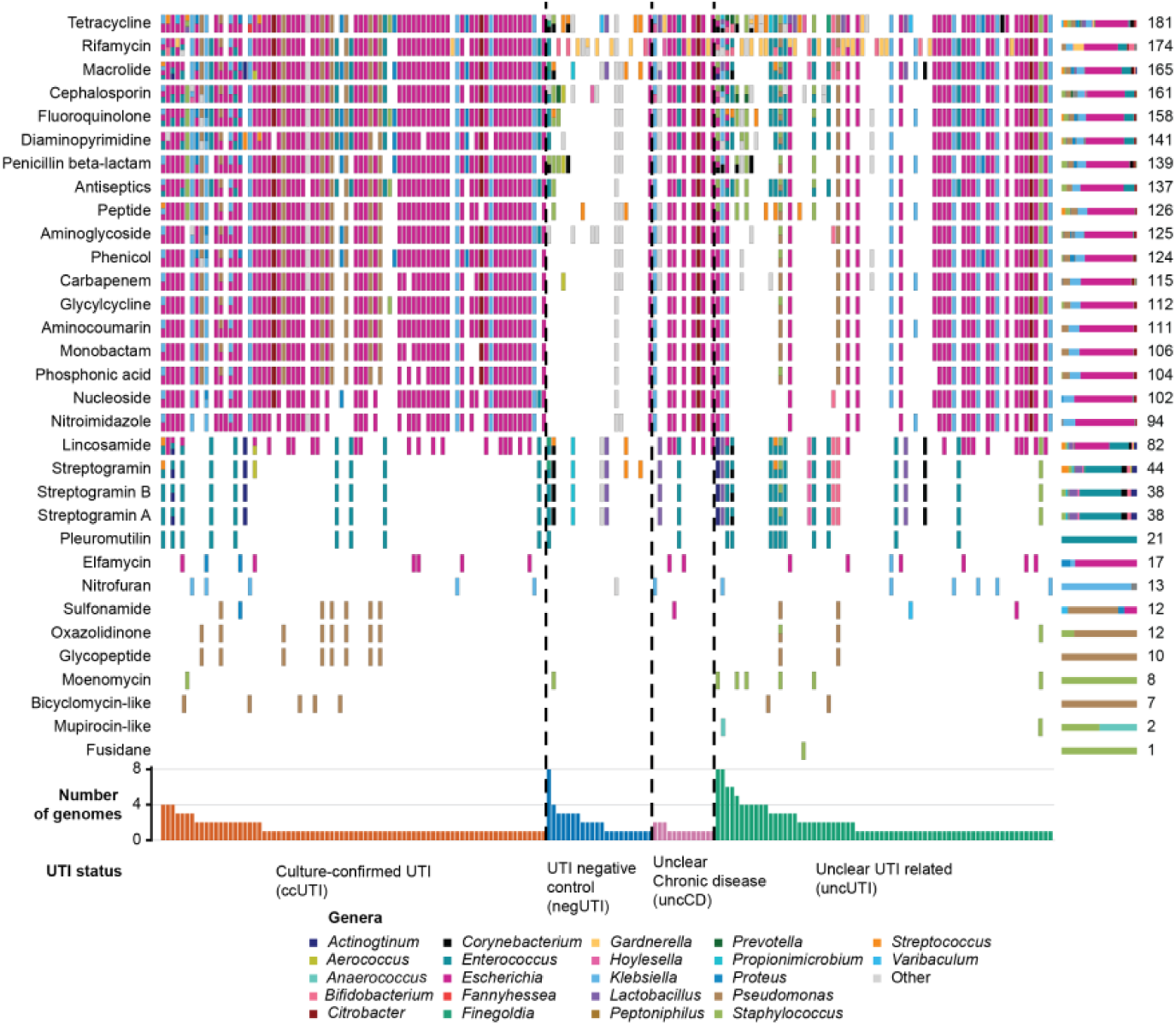
Widespread distribution of AMR across the urobiome. The matrix shows the presence or absence of AMR genes in MAGs recovered from our analysis, organized by samples to highlight co-occurrence of multiple AMR genes. Antibiotics are grouped by drug class, displayed in no particular order. Each cell is a stacked bar with colors indicating the genera containing the AMR genes. On the right, stacked bar plots summarize the proportion of AMR genes per drug class, alongside the total number of AMR genes identified in each drug class. This visualization allows comparison of AMR diversity within and across samples and highlights which bacteria genera contribute most to resistance for each class.

We next sought to determine how AMR genes are distributed across the community. By aggregating the AMR profiles across samples and grouping them by drug class, we found that most of the antibiotic resistances were associated with broad spectrum drugs, including the of rifamycin, tetracycline, macrolide and fluoroquinolone classes (**Fig. 5**). At the mechanistic level, community AMR repertoire was predominated by efflux pump-related mechanisms (70%), with enzymatic resistance accounting for only 25% of the total repertoire. Since efflux pumps act at the individual cell level [95], this suggests that much resistance in these communities is largely mediated through cell-autonomous mechanisms, although these can still be influenced by ecological factors [96,97]. Nevertheless, enzymatic resistance was detected across multiple community members and, given the extracellular mode of action of some (e.g., beta-lactamases), these mechanisms retain the potential to confer community-level antibiotic shielding to neighboring susceptible bacteria [98]. Independently, we observed that multiple genomes carry resistance to the same broad classification of antibiotic class, possibly reflecting Black Queen dynamics [99], and suggesting resistance redundancy within the community, especially observed for *Escherichia* (**Fig. 5**). While the greater abundance of AMR in the ccUTI group could reflect survivor bias of the microbes in patients treated with antibiotics, AMRs were also found in negUTI samples but in a lesser proportion, consistent with previous studies suggesting that microbiomes can act as AMR reservoirs [100,101]. This is further supported by experimental studies and theoretical models, showing that interactions among community members can influence antibiotic efficacy [89,102,103]; for example, by modifying pathogen susceptibility through resistance encoded in commensals [104] or through metabolic cross-feeding that reduces drug effectiveness [105]. These studies combined with our results highlight the importance of considering AMR as a community-level feature of the urobiome.

### Genome-scale analysis reveals phylogenetic and ecological diversity of *E. coli* in the urobiome

*E. coli* inhabits multiple ecological niches in the human body beyond the urinary tract, including the gut [106]. *E. coli* also commonly occurs in the environment and other animal hosts, reflecting its ecological versatility [107]. Given the low phylogenetic diversity of *E. coli* genomes in the urine samples that we described in this manuscript (**Fig. 4C**), we compared them to approximately 50,000 *E. coli* genomes that we obtained from the BV-BRC database [57] encompassing isolates from urine studies and diverse other environments. We then used PopPUNK [59] to cluster the genomes based on whole-genome similarity using k-mer-based distances, detecting 1,916 distinct clusters, reflecting extensive genomic diversity across the strains (**Tab. S5**). For our comparative analyses, we then selected one representative genome per cluster, prioritizing isolates with annotated isolation sources, and combined them with the *E. coli* MAGs recovered in our study (**Fig. 4A**).

To classify all *E. coli* genomes into major phylotypes or phylogroups (A, B1, B2, C, D, E, F, and G), we applied the *in-silico* Clermont phylotyping scheme [108] (**Fig. 6A, 6B**). Consistent with previous reports [109], the majority of UTI-associated genomes belonged to the B2 group [110,111], but genomes from other phylotypes were also recovered from urine samples (**Fig. 6B**). To further assess the relationships between the *E. coli* strains, we constructed a phylogenetic tree using Mash distances, annotated with the detected phylotypes (**Fig. 6A**). The resulting topology recapitulated *E*. *coli* phylotypes, with incongruences likely reflecting differences in sequence coverages. To assess whether genome quality could explain these discordances, we compared the completion and contamination of concordant and discordant phylotype annotations (**Fig. S9**) and found no statistically significant difference between groups, suggesting that assembly quality alone does not account for the observed mismatches. This is consistent with known limitations of phylotype classification methods based on a small number of marker genes and nucleotides variations [112]. The resulting phylogeny is consistent with previous reports that urine-associated *E. coli* encompasses diverse phylogenetic lineages [113]. While B2 remains the predominant UTI-associated phylogroup (**Fig. 6B**), the presence of diverse *E. coli* lineages across UTI status highlights the ecological plasticity of the species [115].

**Figure 6.**
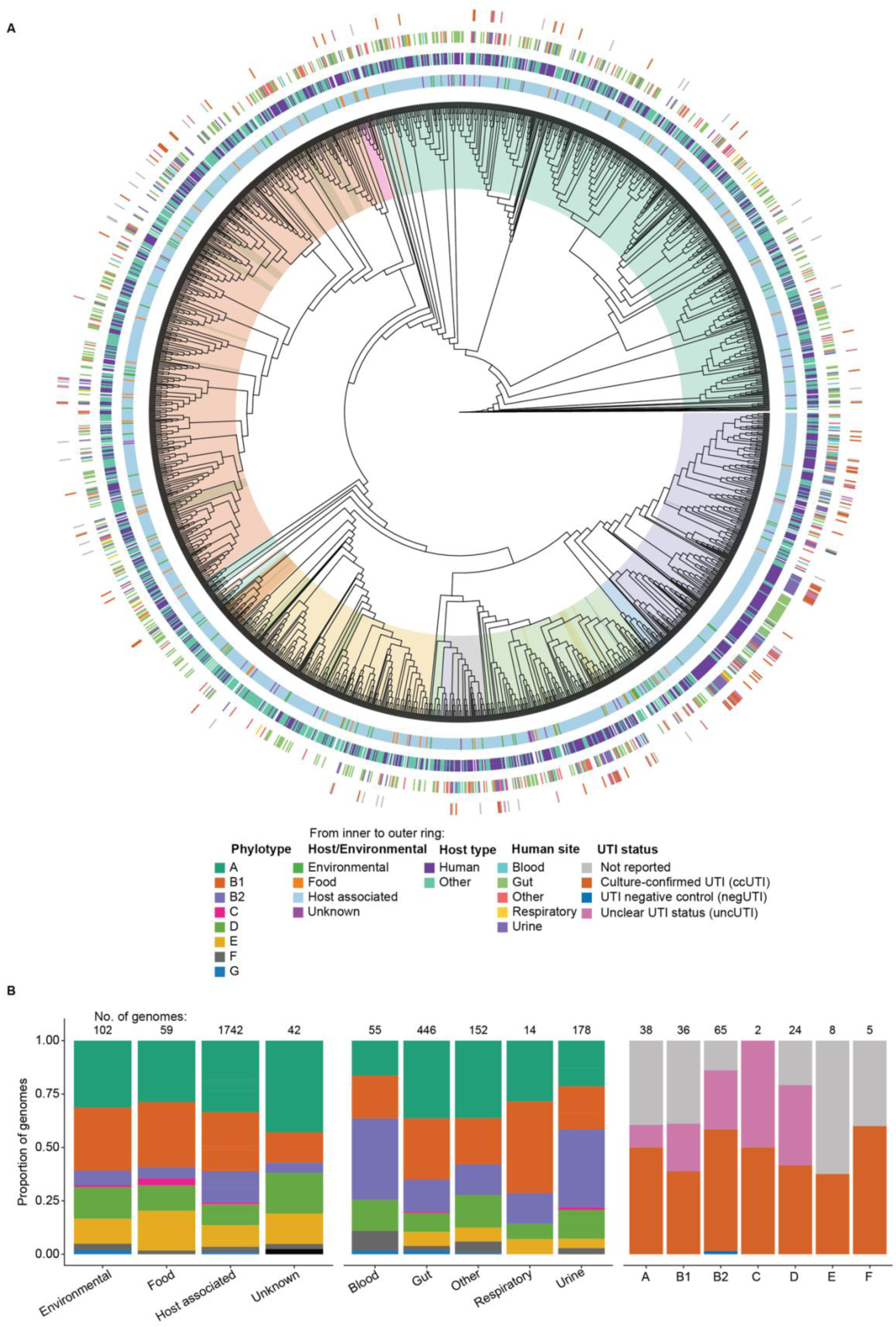
*Escherichia coli* lineages span multiple phylotypes and ecological niches. **A)** Phylogenetic tree including both reference *E. coli* genomes and genomes recovered from our analysis. Branch lengths are not drawn according to scale, as the *E. coli* phylogeny is highly divergent, and large-scale branches hamper the visualization. The area behind branches is shadowed by phylotype, concentric rings display associated metadata and, isolation source from inner to outer ring: host or environmental related, host type, human site, and UTI status at the time of sampling. *E. coli* genomes are phylogenetically and ecologically diverse, with phylotypes largely consistent with the phylogeny and lineages derived from multiple isolation sources. **B)** Overview of the dataset composition. The number of genomes per category is listed above the bars: the first plot (left) shows all isolation sources colored by phylotype, the middle plot shows specific human sites (subset of “host associated”) also colored by phylotype, and the right plot shows urine-derived genomes colored by UTI status.

### Phylotype-specific genomic signatures dominate the functional landscape of urine *Escherichia coli*

To investigate whether *E. coli* lineages exhibit genomic features linked to adaptation to specific niches such as the urinary tract, we constructed a pangenome with the representatives’ dataset using Panaroo [61] (**Fig. 7A**). We defined genes present in ≥95% of *E. coli* genomes as core (2,480 genes), those present in 15-95% as shell (3,091 genes), and those present in <15% as accessory or cloud (59,683 genes). Heaps’ law (α = 0.58) indicates that *E. coli* has an open pangenome (**Fig. 7B**), consistent with previous studies [116–118]. Similarly, when we limited our results to only those genomes labeled as ccUTI, we also observed an open pangenome (α = 0.59, **Fig. S10**). While the numbers of core and shell genes are broadly similar when compared with previous studies (core: 2,000 - 3,000 genes, shell: 3,000 - 5,000 genes) [119,120], the size of the accessory genome varies considerably, ranging from 10,000 to 160,000 genes [117,121], likely reflecting differences in the number, diversity, and quality of genomes included in each analysis.

**Figure 7.**
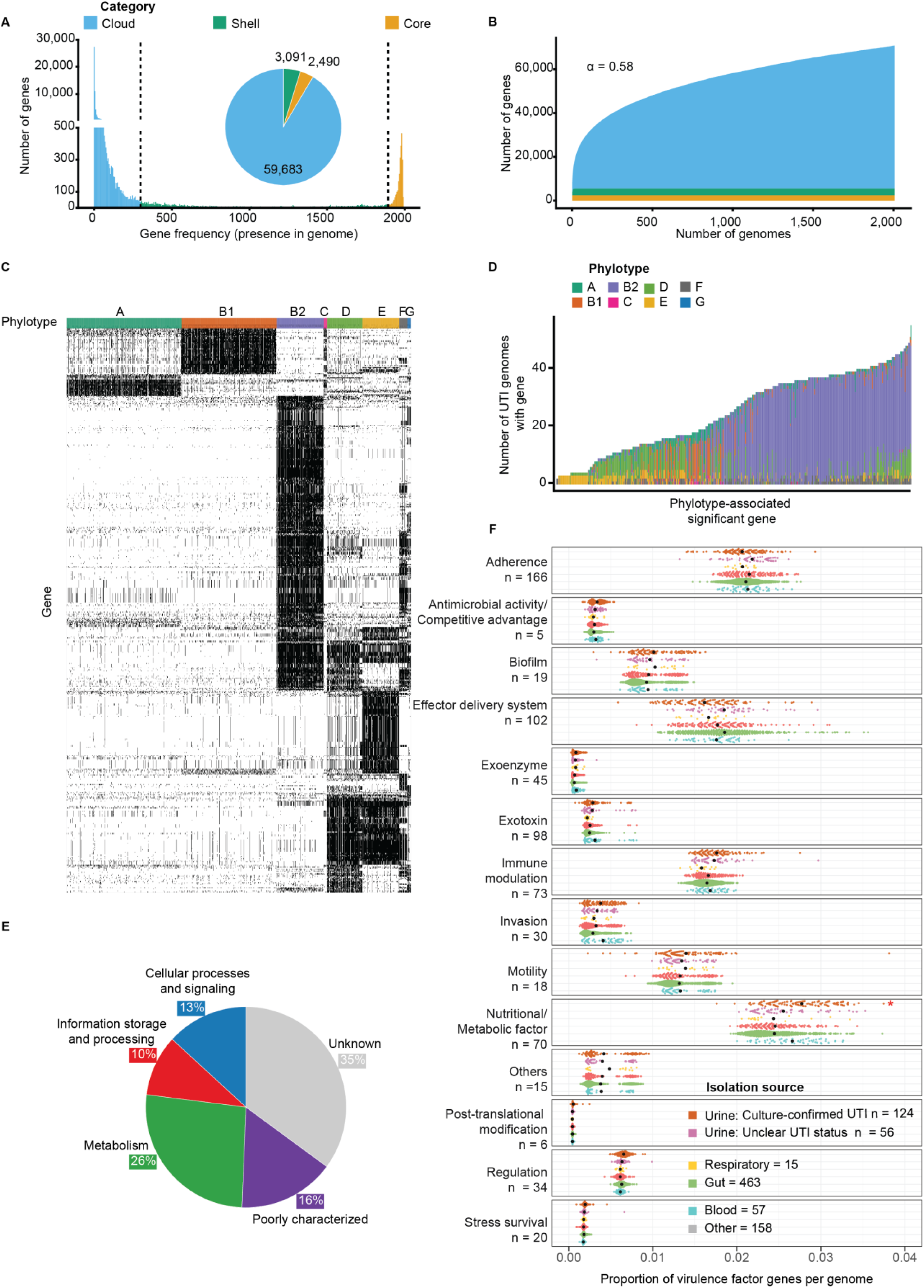
High genomic diversity of *E. coli* with limited association to UTI status. **A)** Distribution of core, shell, and cloud genes in the pangenome of *E. coli*, showing the number of genes in each category. **B)** Pangenome growth curve based on Heaps’ law (α = 0.58), indicating an open pangenome. Core and shell gene numbers stabilize, whereas cloud/accessory genes continue to increase with additional genomes. **C)** Heatmap of presence–absence for genes significantly associated with phylotypes identified in the pan-GWAS with Scoary (p value < 0.05, % sensitivity 75 % and specificity 75%, fisher’s exact test and Bonferroni correction for multiple testing). **D)** Pie plot displays the proportion of the statistically significant genes in panel C) genes grouped by functional category (COG class). **E)** Distribution of phylotype-associated genes in UTI-positive genomes, illustrating overlap between significant genes found in panel C among UTI-positive groups of genomes. **F)** Proportion of VF genes per genome, stratified by VF category and isolation source. Each point represents an individual genome, and its correspondent value is the within-genome proportion of genes assigned to a given VF category (numbers below correspond to the number of genes per category). Points are colored by isolation source. Black dots indicated the median proportion for each group, and numbers below each isolation source indicate the number of genomes included. Statistical differences between isolation sources were assessed using Kruskal-Wallis tests followed by Dunn’s post hoc test with Bonferroni correction for multiple testing and filtered. Only results with adjusted p < 0.05 were considered significant. The nutritional /metabolic VF category was significantly enriched in urine isolated within the ccUTI group compared to the other isolation sources.

To identify gene families that are significantly associated with isolation source, phylotype, or UTI status, we performed a pan-GWAS analysis with Scoary [62]. Across all traits, 550 genes were significantly associated (p value < 0.05, % sensitivity 75 % and % specificity 75%, Fisher’s exact test and Bonferroni correction for multiple testing, **Fig. 7C**, **Fig. 7D**). Interestingly, all significant associations solely corresponded to phylotype, whereas no gene family was significantly linked to isolation source or clinical phenotype, consistent with previous studies showing stronger genomic signatures at the phylogroup level than by niche [117,122]. Among phylotypes, the B2 group showed the highest enrichment of gene families, particularly those involved in metabolism and nutrient transport, consistent with previous reports highlighting conserved metabolic traits in B2 genomes (**Fig. S11**) [123,124]. To explore whether phylotype-associated genes can also be associated with UTI development, we examined their distribution among ccUTI genomes. Although not statistically associated with UTI status, most gene families present in UTI-derived genomes predominantly belonged to the B2 phylotype (**Fig 6B**, **Fig. 7E**).

Next, we investigated functional genomic traits to understand whether the urinary niche leads to potentially specific pathogenic signatures in *E. coli.* We analyzed the virulence factor (VF) repertoire across genomes the representative dataset of genomes using the VFDB database (**Fig. 7F**) [66], and quantified for each genome individually the proportion of abundance of each VF category relative to the total number of genes annotated in that genome. After grouping genomes by isolation source, we compared the distribution of VF category proportions across body sites and tested statistically significant differences. Of all VF categories examined, only nutritional/metabolic factors showed a significant enrichment in ccUTI isolates (**Fig. 7F**). Urinary *E. coli* has previously been described to rely on essential metabolic pathways for growth [125], including amino acid transport and metabolism [126], and the utilization of alternative carbon sources [127]. Rather than reflecting a single, specialized lineage, our results indicate an enrichment of metabolic factors likely reflecting ecological constraints imposed by the host and the surrounding urobiome, including interactions such as metabolic cross-feeding [128], and modulation of enzyme activity [129]. Consequently, even within a single phylotype, *E. coli* strains can acquire or maintain traits that support fitness within the urinary tract, as observed in strains consequently persisting in both intestinal and urinary tracts depending on the context [130] suggesting an opportunistic model of infection.

## CONCLUSIONS

UTIs impact tens of millions of women per year, and to a lesser extent men. Despite its clinical importance, the composition and functional capabilities of the resident microbiome of the urinary tract remain largely unexplored. Our study highlights the interplay between community structure, functional potential, and lineage-specific traits in shaping the dynamics of the urobiome. We profiled the urobiome of individuals with and without UTI at the time of sampling, identifying distinct clusters that are not strictly associated with the presence or absence of a UTI. Our analyses revealed broad phylogenetic diversity across the urobiome, with uropathobiont genera showing closer phylogenetic relationships to each other than to vaginal and skin associated bacteria. We also highlighted widespread AMR potential, but limited adaptation of the functional potential of urine-associated *E. coli*. While considerations such as variability across metagenomic datasets, potential MAG contamination, and incomplete assemblies may affect the annotation and interpretation of functional and resistance profiles, our findings suggest that uropathobiont success in the urinary tract may be shaped not only by intrinsic metabolic advantages but also by interactions with the surrounding urobiome. Future work will therefore need to explore how host environmental factors, community interactions, and metabolism shape uropathogen survival and infection dynamics. Integrating this ecological context will be essential to better understand infection processes, and to develop alternative treatments to reduce the prevalence of UTIs.

## DECLARATIONS

### Ethics approval and consent to participate

Not applicable.

### Consent for publication

Not applicable.

### Availability of data and material

The sequencing data used in this study can be found in the following publicly available sources: NCBI (BioProjects IDs: PRJNA700071, PRJNA679884, PRJNA385350, PRJNA801448, PRJNA400628 and PRJNA680735) and ENA (Project accession: PRJEB45363).

### Author contributions

D.G.V., J.H.H.M.W., B.E.D., and M.F.S. conceptualized and designed the study. D.G.V. performed the experiments and collected the data. D.G.V. analyzed data. D.G.V., B.E.D., and M.F.S drafted the manuscript, and C.S., A.J.W., M.V., and J.H.H.M.W. revised the initial draft. All authors approved the final manuscript.

## Acknowledgments

We would like to thank the Urinary tract infections revisited (UTIr) consortium for the fruitful discussion and feedback. We thank Jan Kees van Amerongen for the invaluable support and the maintenance of the computational infrastructure.

## Additional files

Additional file 1. Additional_file_1.xls. Supplementary Tables.

Supplementary Table 1. Summary of samples included in the metagenomic analyses, including sequencing methodology and available sample-level metadata as provided by the original studies (n = 433).

Supplementary Table 2. Overview of the sample selection criteria for all studies included in this work. Supplementary Table 3. Quality and taxonomical data for the MAGs recovered in this study.

Supplementary Table 4. An overview of antimicrobial resistance (AMR) profiles identified in metagenome-assembled genomes (MAGs), including CARD-based annotations in antibiotic class, resistance type, and quality control metrics.

Supplementary Table 5. Information on the *E*. *coli* isolates used in this work, including their predicted phylotype.

Additional file 2. Additional_file_2.docx. Supplementary Figures.

Supplementary Figure 1. The number of paired reads for pre- and post-processing across studies varies. Comparison of the number of reads obtained from every study before (raw) and after quality control (QC), box plot shows the distribution of the read cunts.

Supplementary Figure 2. Taxonomic classification is consistent across studies. Percentage of paired reads classified by Metaphlan per sample is shown, grouped by their original study.

Supplementary Figure 3. PCoA reveals no clustering for UTI status or study. Principal coordinate analysis (PCoA) based on Euclidean distances of the transformed relative abundances, colored by A) study and B) UTI status.

Supplementary Figure 4. Richness and alpha diversity positive trends through age. Distribution of the alpha diversity through age, divided by UTI status. The linear model was fitted to the alpha diversity of the samples and correlated with the age of every sample. Positive correlation was observed for the UTI negative group and the alpha diversity characterized by an increase of it through the age.

Supplementary Figure 5. Differential abundance by age. A) Differential abundance of bacterial species across age identified using MaAslin3. B) Differential prevalence of bacterial species across age, shown as log fold change (odds ratio).

Supplementary Figure 6. The number of low, medium, and high-quality MAGs were successfully recovered across all metagenomes. High-quality genomes were recovered despite the low microbial biomass in many urine samples.

Supplementary Figure 7. Bacteria commonly identified as pathogenic are clade-specific, while other taxa are phylogenetically diverse. Distribution of phylogenetic distances within the main genera shown in Fig. 4C. Boxplots represent the range and variation of intra-genus phylogenetic distances, highlighting differences in lineage diversity across genera. Clinical relevance was determined based on clinical reports of most-prevalent bacteria causing UTI.

Supplementary Figure 8. Prevalence of predicted pathogenic genera across UTI status groups. Bar plot showing the prevalence of genera across all samples, grouped by predicted pathogenicity based on PathogenFinder2 predictions. Genera are categorized as predicted human pathogens or non-pathogens. Non-common uropathobionts presented pathogenic potential in the negUTI group.

Supplemental Figure 9. Genome quality does not explain phylotype discordances. Violin plots showing completeness and contamination of E. coli MAGs grouped by concordance between ClermonTyping-assigned phylotypes and phylogenetic placement on the whole-genome tree. No significant differences were observed between concordant and discordant groups (Wilcoxon test, p > 0.05).

Supplementary Figure 10. UTI-associated E. coli has an open pangenome. Pangenome growth curve basedon Heaps’ law (α = 0.59) for only UTI positive *E. coli* genomes, indicating an open pangenome. Core and shell gene numbers stabilize, whereas cloud/accessory genes continue to increase with additional genomes being added.

Supplementary Figure 11. Genes detected by pan-GWAS to be significantly associated with phylotypes are related to various COG categories. Heatmap of presence–absence for genes significantly associated with phylotypes identified in the pan-GWAS, colored by COG broad category.

## SUPPLEMENTARY FIGURES

**Supplementary Figure 1.**
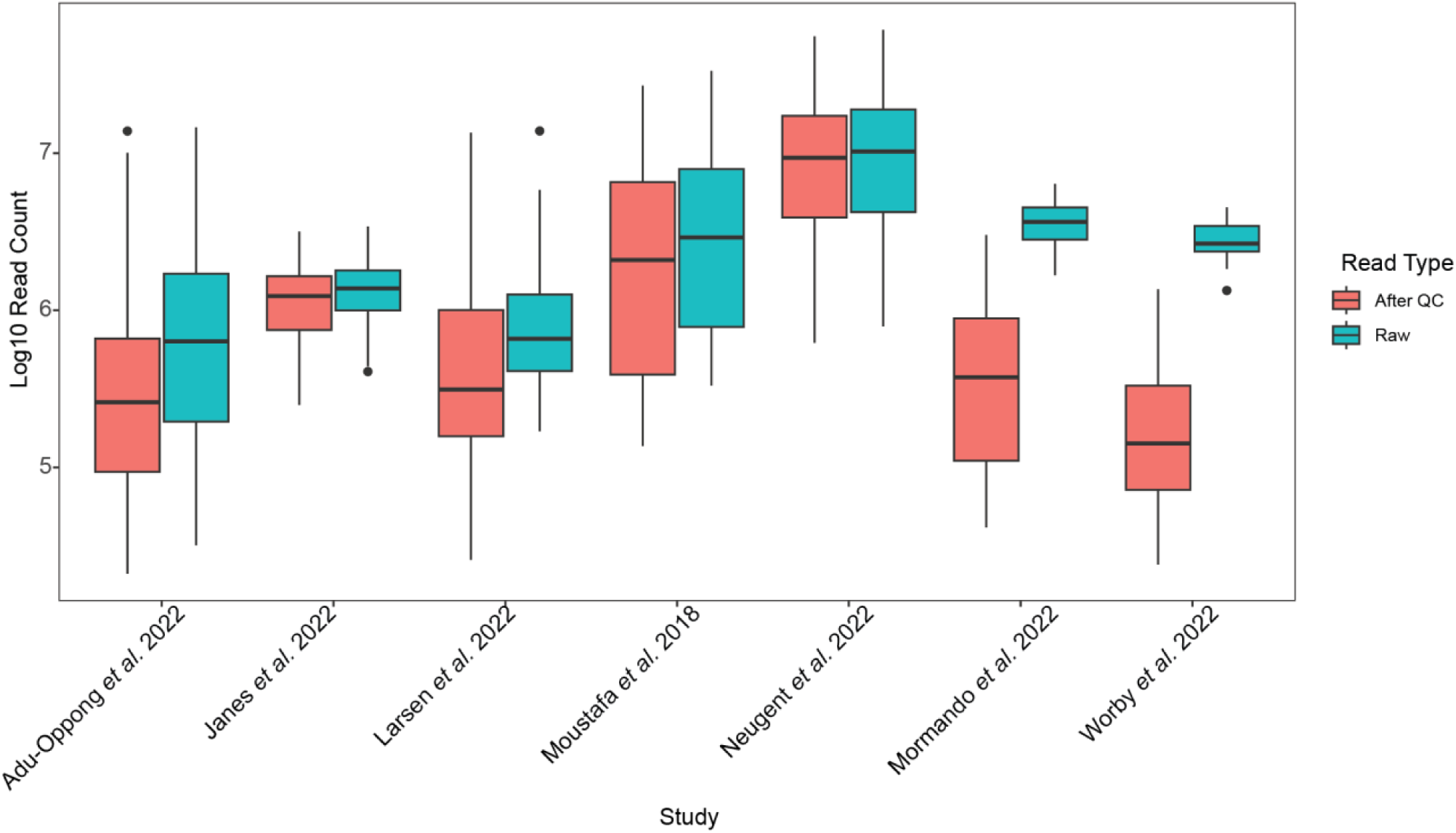
The number of paired reads for pre- and post-processing across studies varies. Comparison of the number of paired reads obtained from every study before (raw) and after quality control (QC), box plot shows the distribution of the read counts.

**Supplementary Figure 2.**
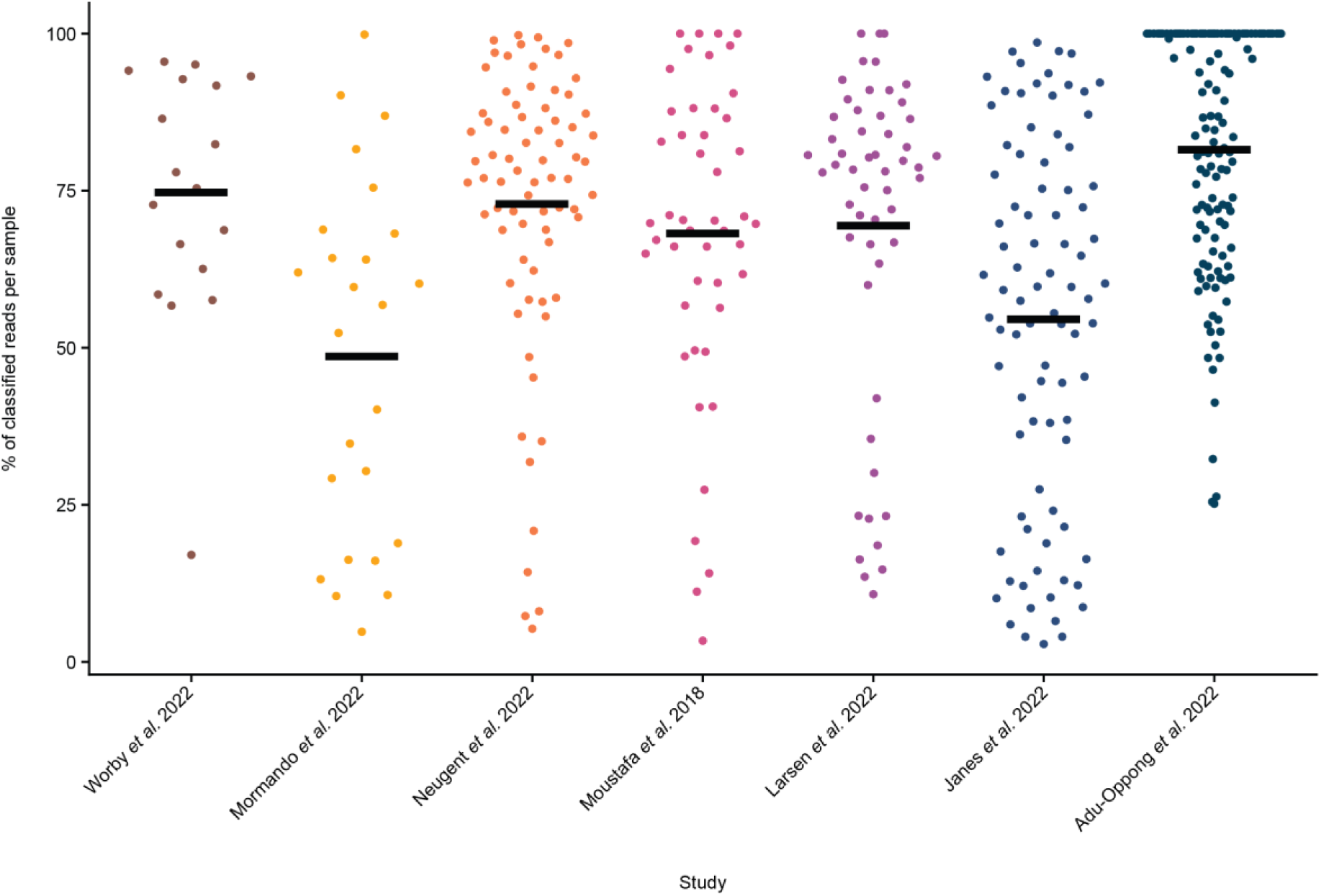
Taxonomic classification is consistent across studies. Percentage of paired reads classified by Metaphlan per sample is shown, grouped by their original study.

**Supplementary Figure 3.**
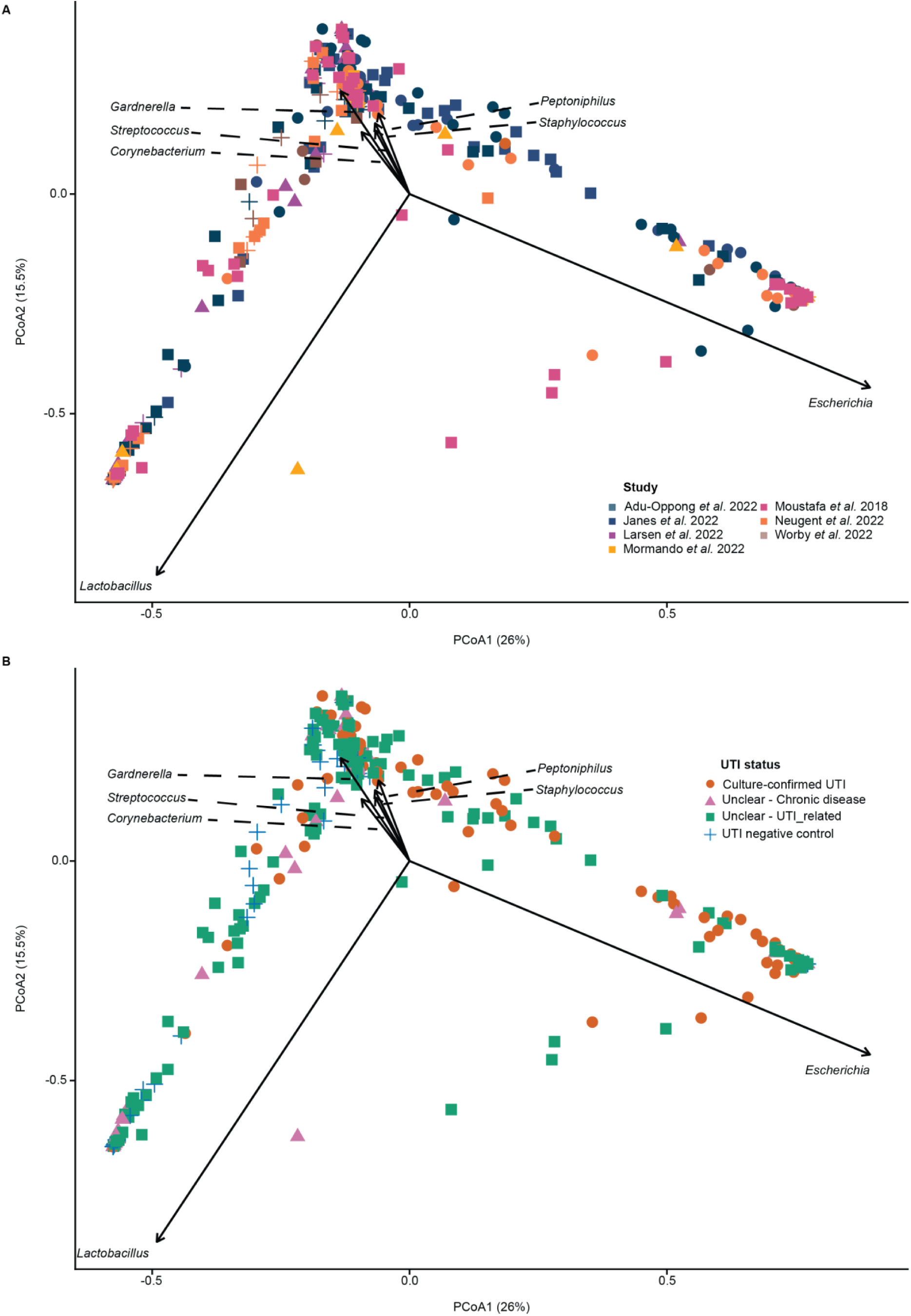
PCoA reveals no clustering for UTI status or study. Principal coordinate analysis (PCoA) based on Euclidean distances of the transformed relative abundances, colored by A) study and B) UTI status.

**Supplementary Figure 4.**
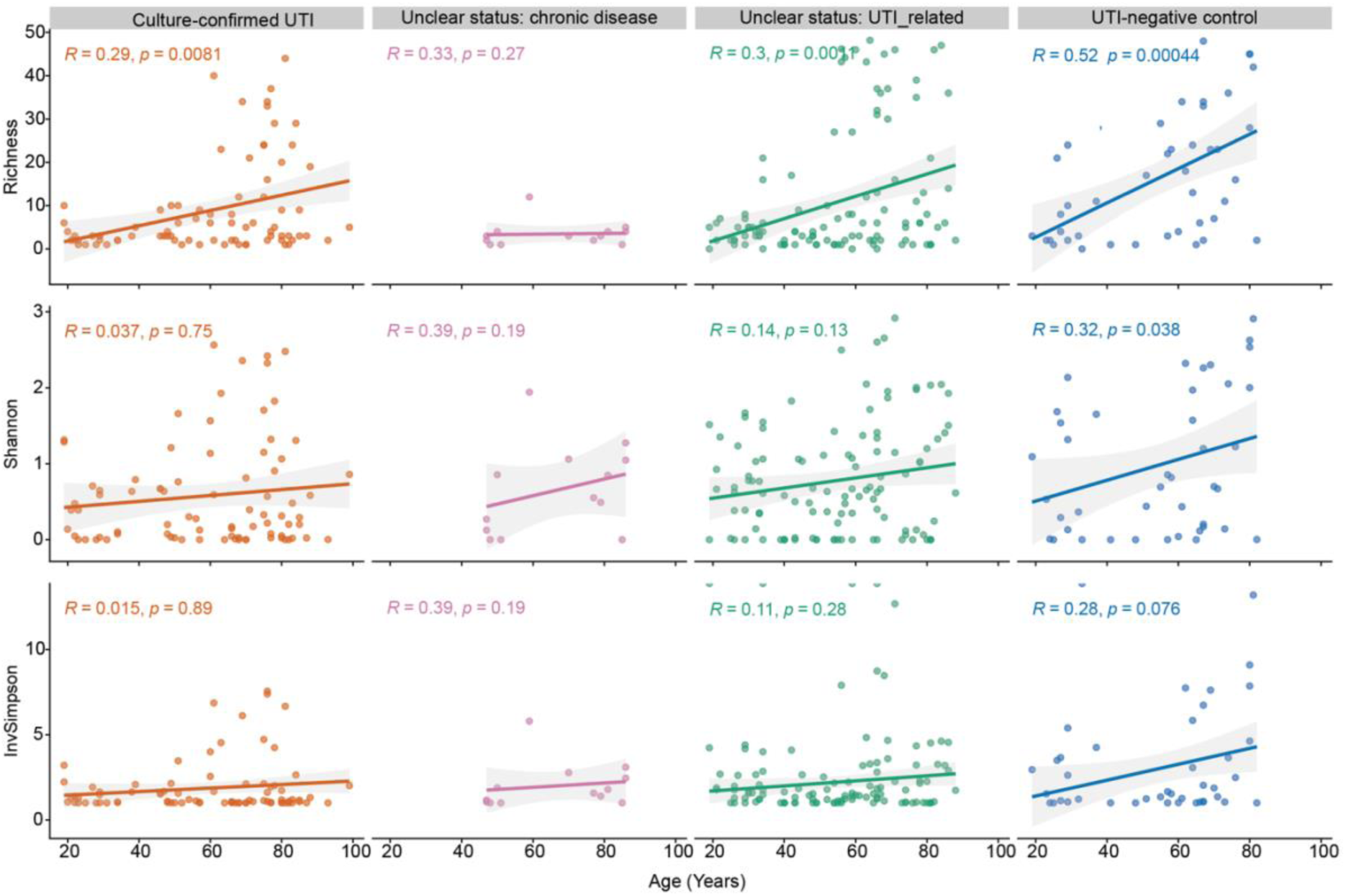
Richness and alpha diversity positive trends through age. Distribution of the alpha diversity through age, divided by UTI status. The linear model was fitted to the alpha diversity of the samples and correlated with the age of every sample. Positive correlation was observed for the UTI negative group and the alpha diversity characterized by an increase of it through the age.

**Supplementary Figure 5.**
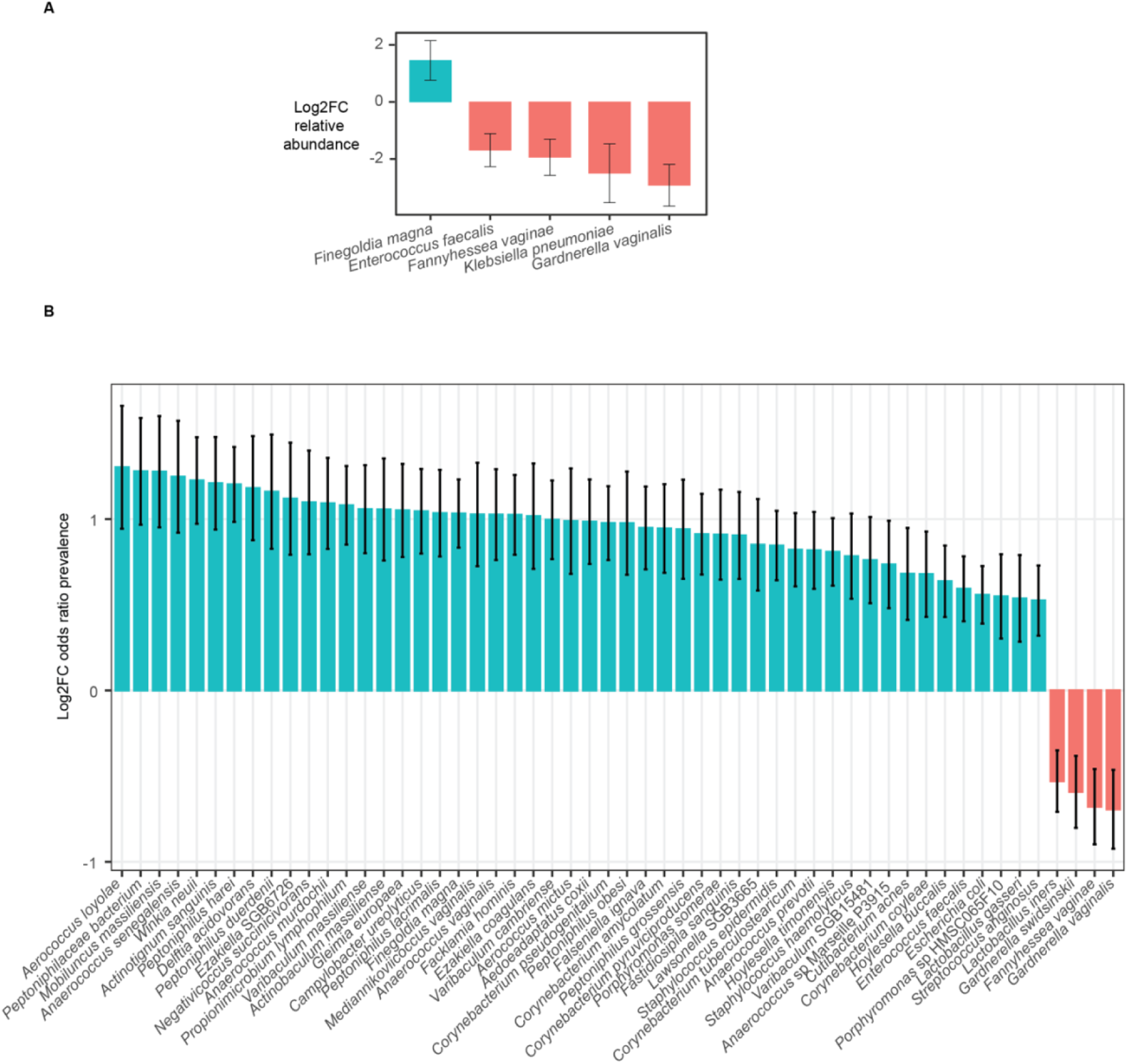
Differential abundance by age. **A)** Differential abundance of bacterial species across age identified using MaAslin3. **B)** Differential prevalence of bacterial species across age, shown as log fold change (odds ratio).

**Supplementary Figure 6.**
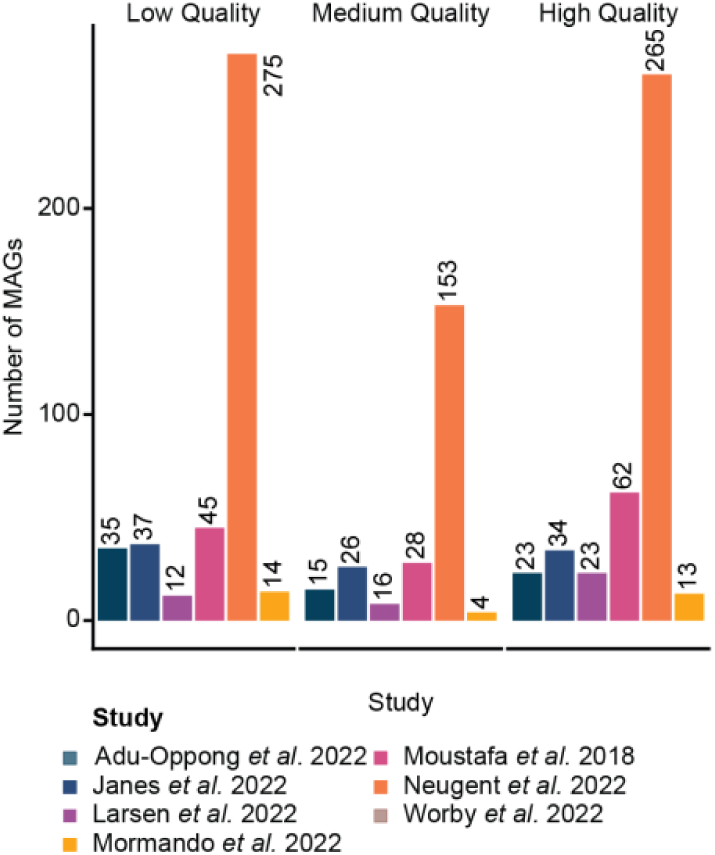
The number of low, medium, and high-quality MAGs were successfully recovered across all metagenomes. High-quality genomes were recovered despite the low microbial biomass in many urine samples.

**Supplementary Figure 7.**
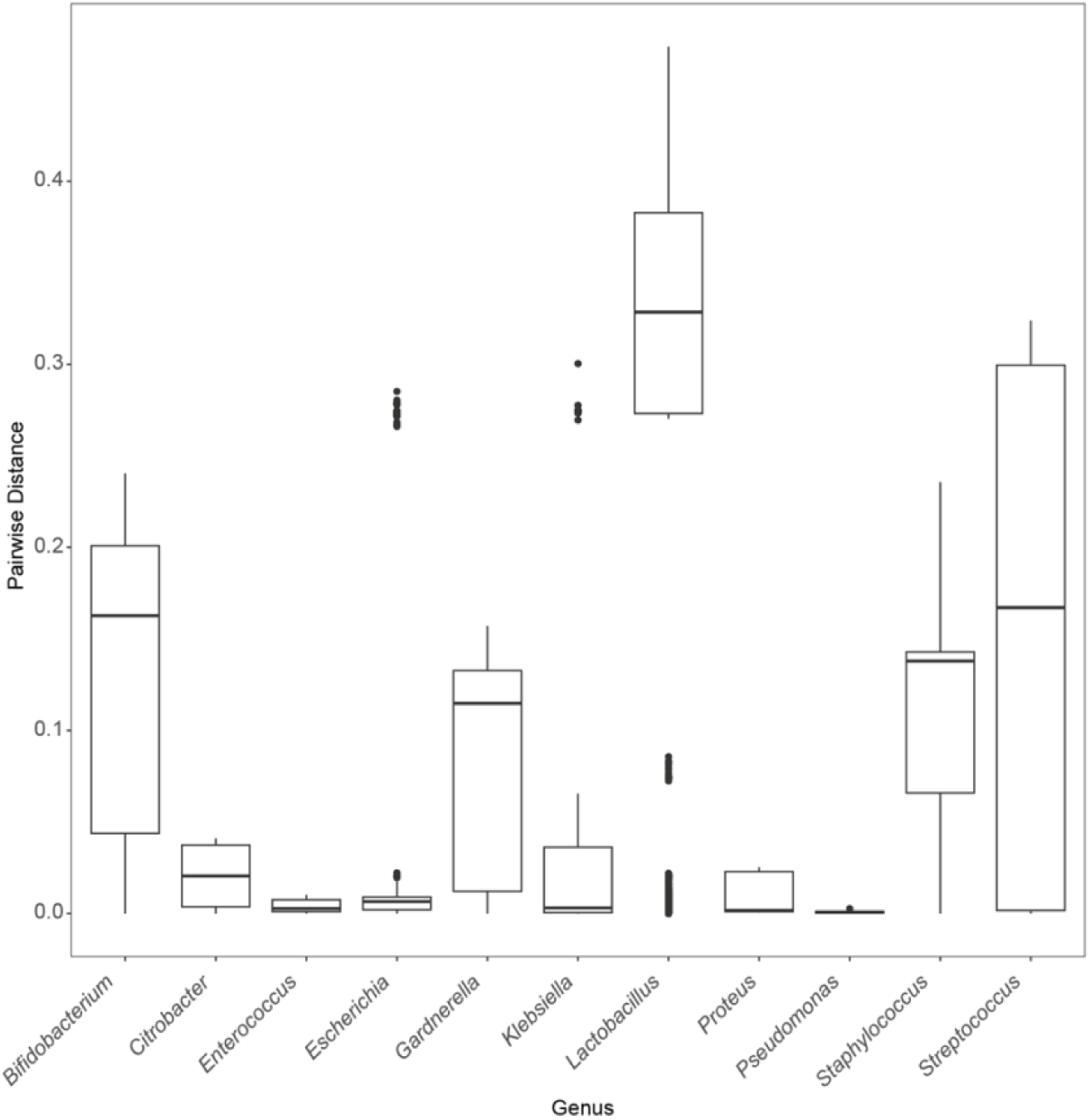
Bacteria commonly identified as pathogenic are clade-specific, while other taxa are phylogenetically diverse. Distribution of phylogenetic distances within the main genera shown in Fig. 4C. Boxplots represent the range and variation of intra-genus phylogenetic distances, highlighting differences in lineage diversity across genera. Clinical relevance was determined based on clinical reports of most-prevalent bacteria causing UTI.

**Supplementary Figure 8.**
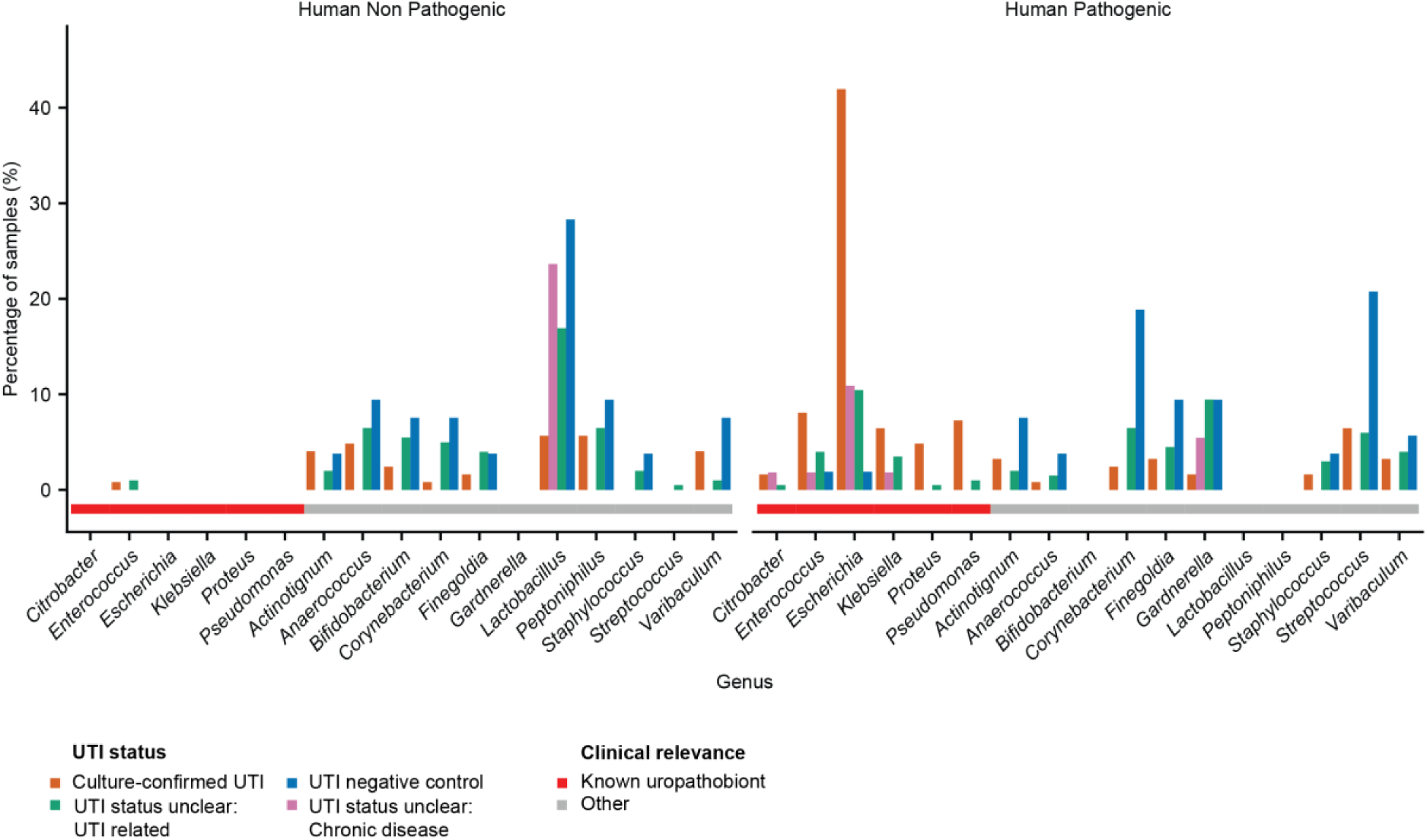
**Prevalence of predicted pathogenic genera across UTI status groups**. Bar plot showing the prevalence of genera across all samples, grouped by predicted pathogenicity based on PathogenFinder2 [84] predictions. Genera are categorized as predicted human pathogens or non-pathogens. Non-common uropathobionts presented pathogenic potential in the negUTI group.

**Supplemental Figure 9.**
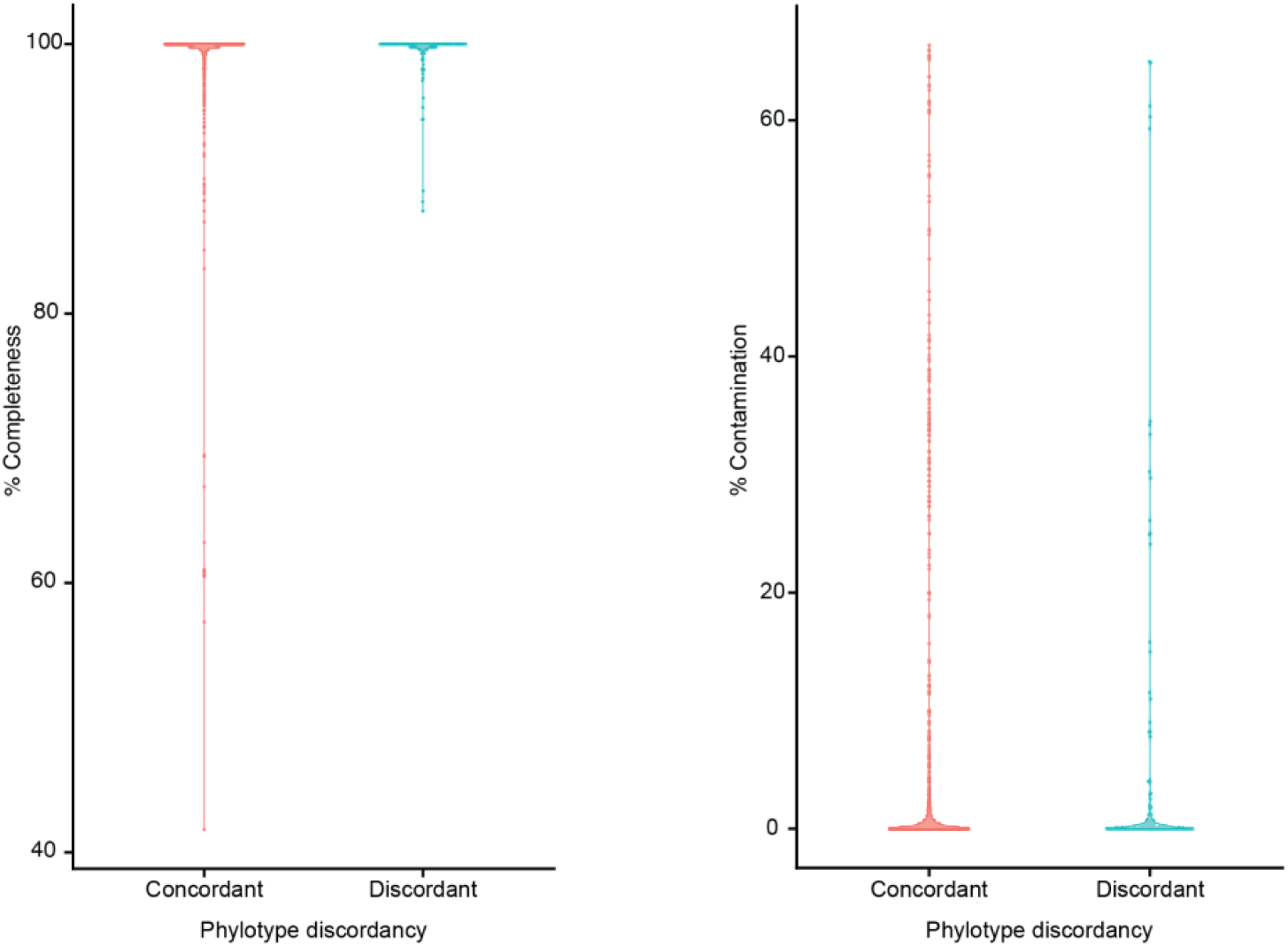
**Genome quality does not explain phylotype discordances**. Violin plots showing completeness and contamination of *E. coli* MAGs grouped by concordance between ClermonTyping-assigned phylotypes and phylogenetic placement on the whole-genome tree. No significant differences were observed between concordant and discordant groups (Wilcoxon test, p > 0.05).

**Supplementary Figure 10.**
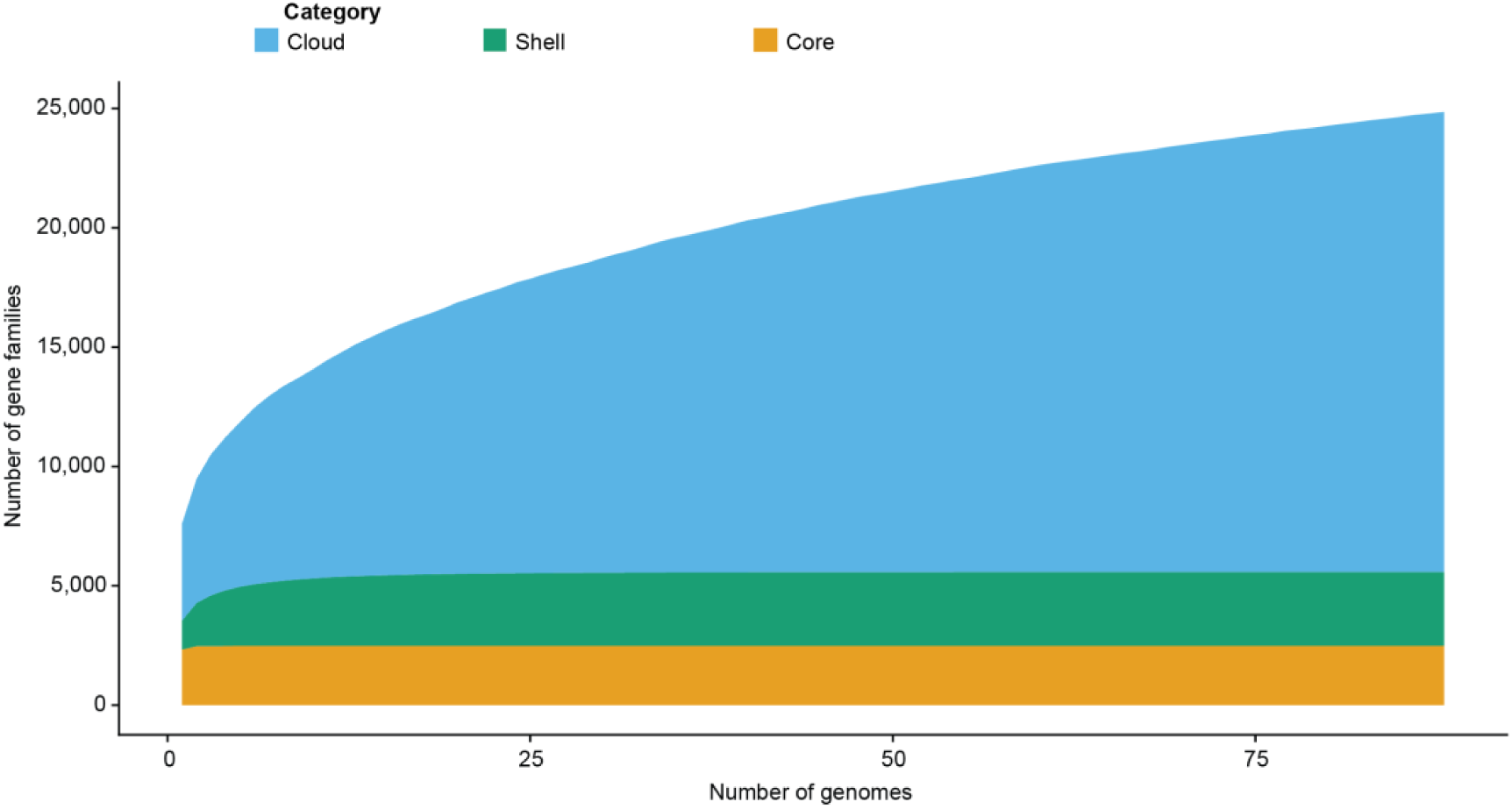
UTI-associated *E. coli* has an open pangenome. Pangenome growth curve based on Heaps’ law (α = 0.59) for only UTI positive *E. coli* genomes, indicating an open pangenome. Core and shell gene numbers stabilize, whereas cloud/accessory genes continue to increase with additional genomes being added.

**Supplementary Figure 11.**
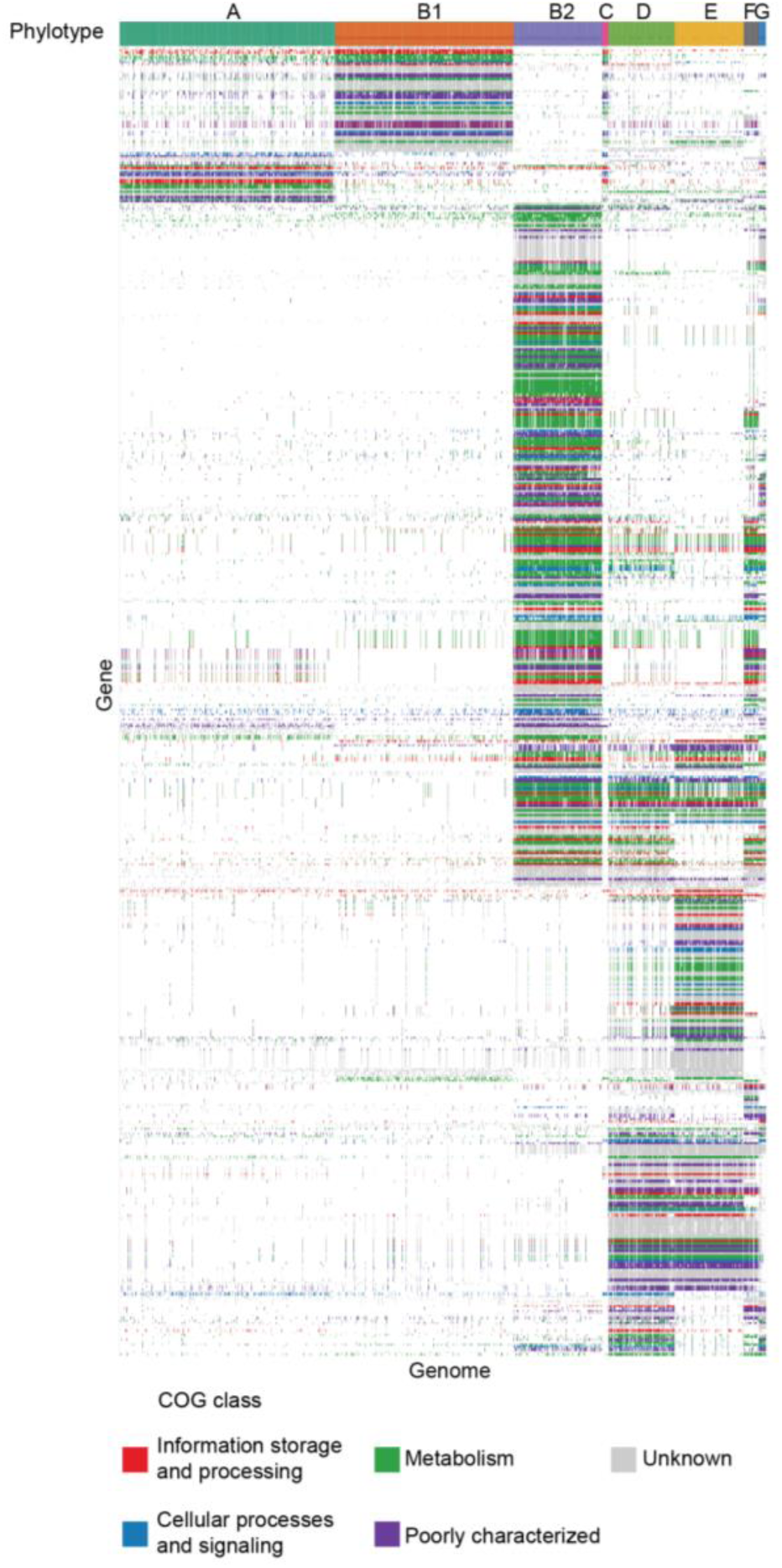
Genes detected by pan-GWAS to be significantly associated with phylotypes are related to various COG categories. Heatmap of presence–absence for genes significantly associated with phylotypes identified in the pan-GWAS, colored by COG broad category.

