## Additional file 2 for "Cross-study metagenomics analysis reveals distinct microbial signatures of urinary tract infections"

### SUPPLEMENTARY FIGURES

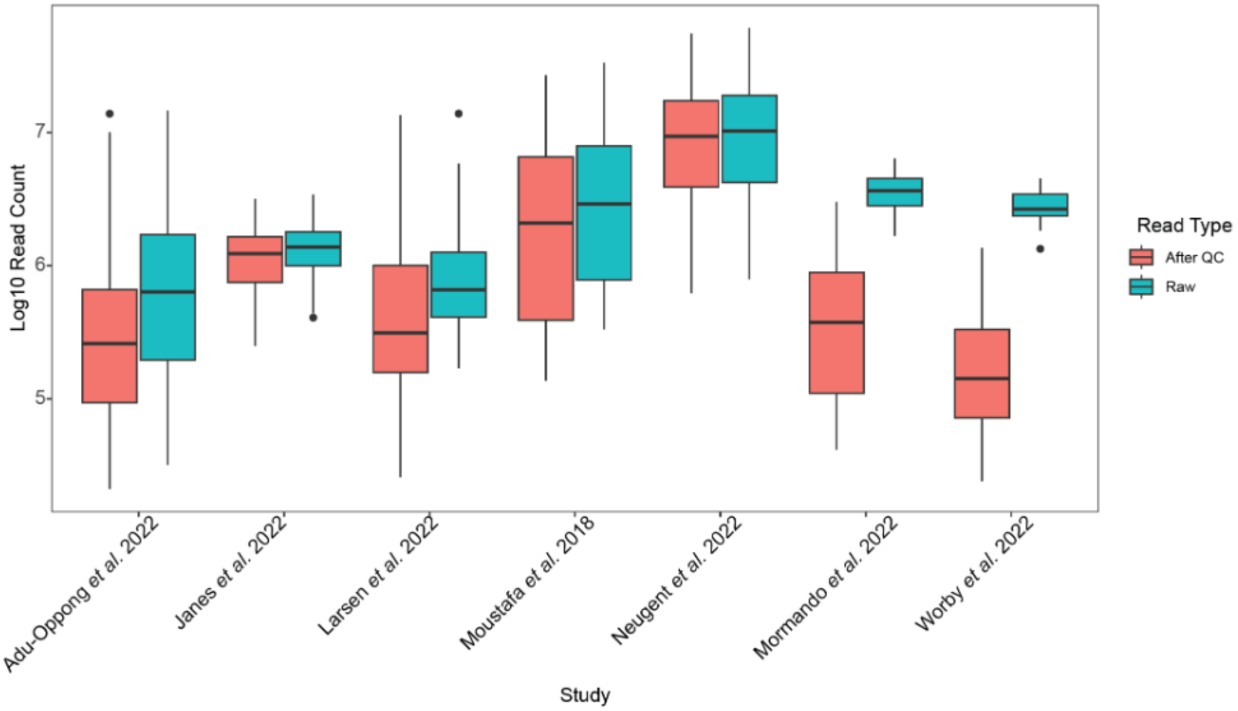


**Supplementary Figure 1. The number of paired reads for pre- and post-processing across studies varies.** Comparison of the number of paired reads obtained from every study before (raw) and after quality control (QC), box plot shows the distribution of the read counts.

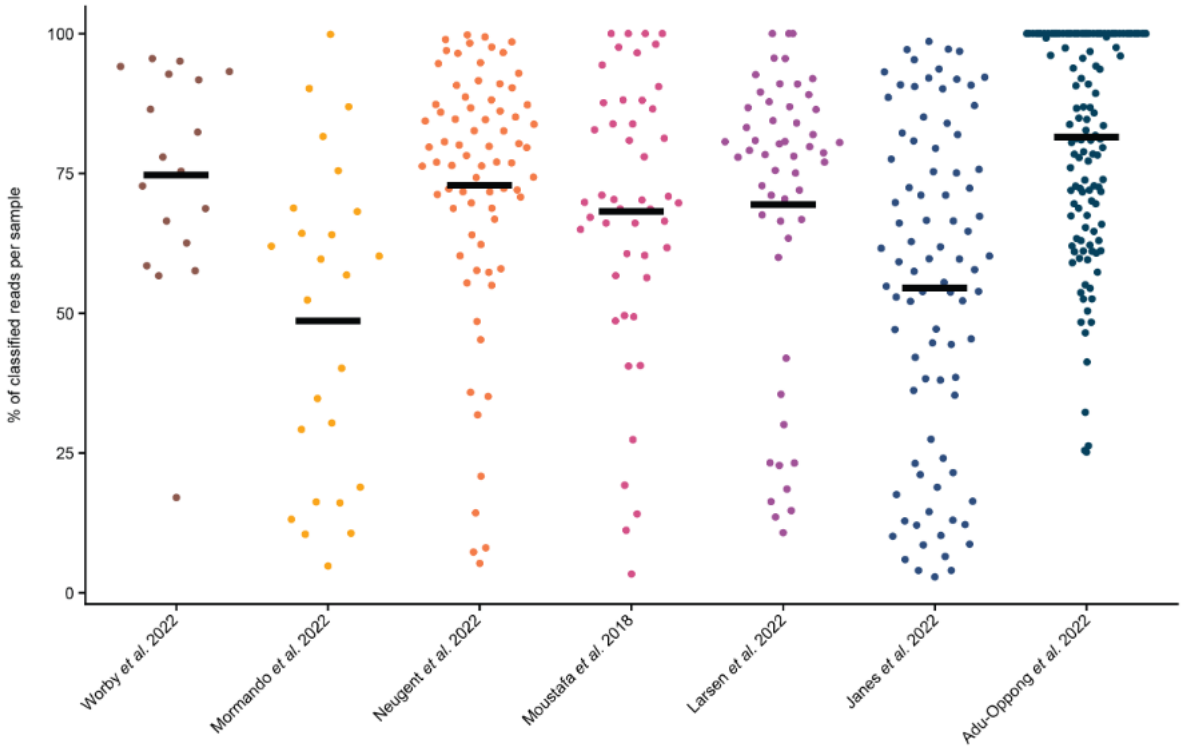


**Supplementary Figure 2**. **Taxonomic classification is consistent across studies.** Percentage of paired reads classified by Metaphlan per sample is shown, grouped by their original study.

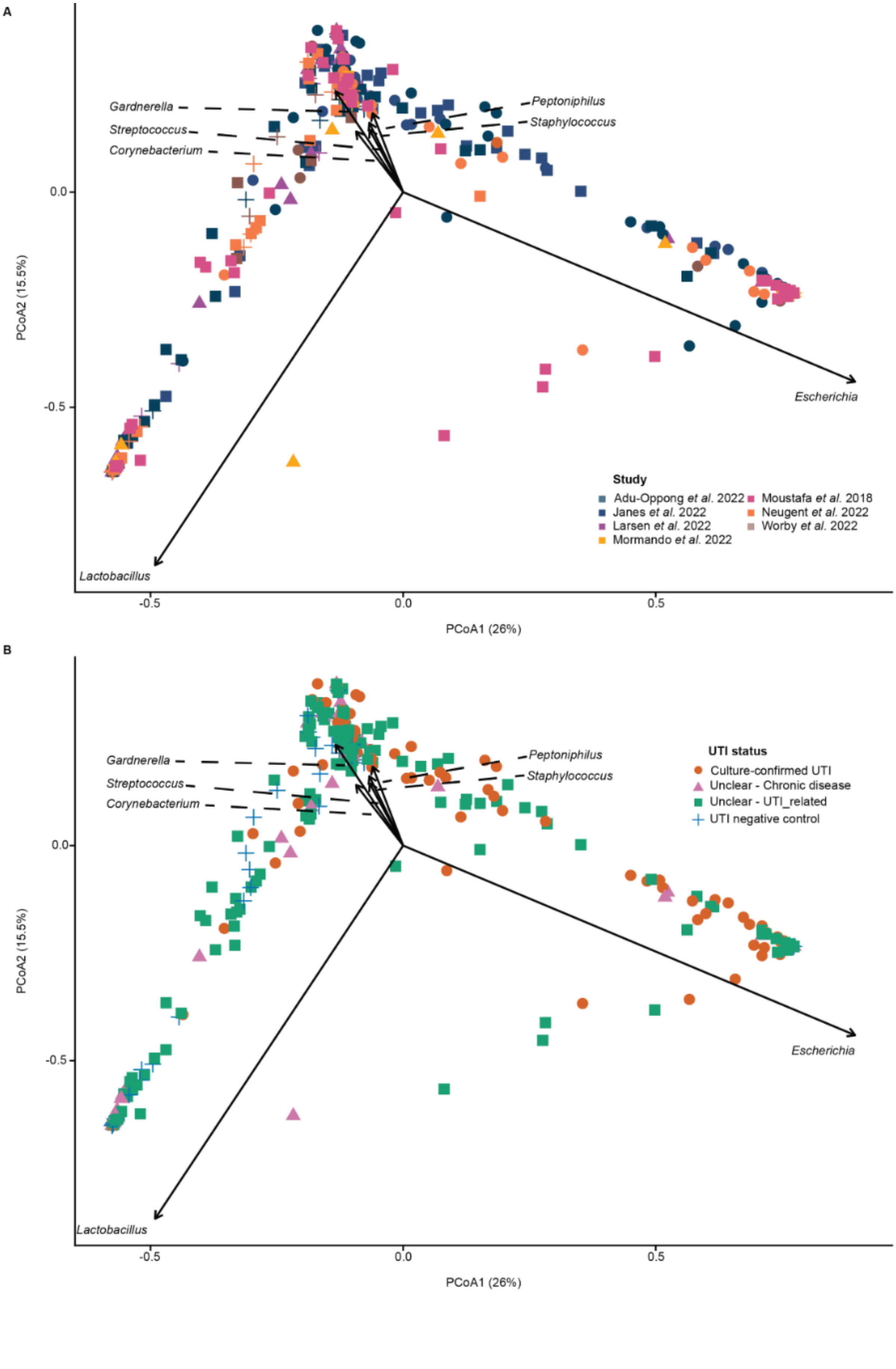

A
Gardnerella
Streptococcus
Corynebacterium
Peptoniphilus
Staphylococcus
Escherichia
Lactobacillus
PCoA2 (15.5%)
PCoA1 (26%)
0.0
-0.5
0.5
Study
Adu-Oppong et al. 2022
Janes et al. 2022
Larsen et al. 2022
Mormando et al. 2022
Moustafa et al. 2018
Neugent et al. 2022
Worby et al. 2022
B
UTI status
Culture-confirmed UTI
Unclear - Chronic disease
Unclear - UTI_related
UTI negative control

**Supplementary Figure 3. PCoA reveals no clustering for UTI status or study.** Principal coordinate analysis (PCoA) based on Euclidean distances of the transformed relative abundances, colored by A) study and B) UTI status.

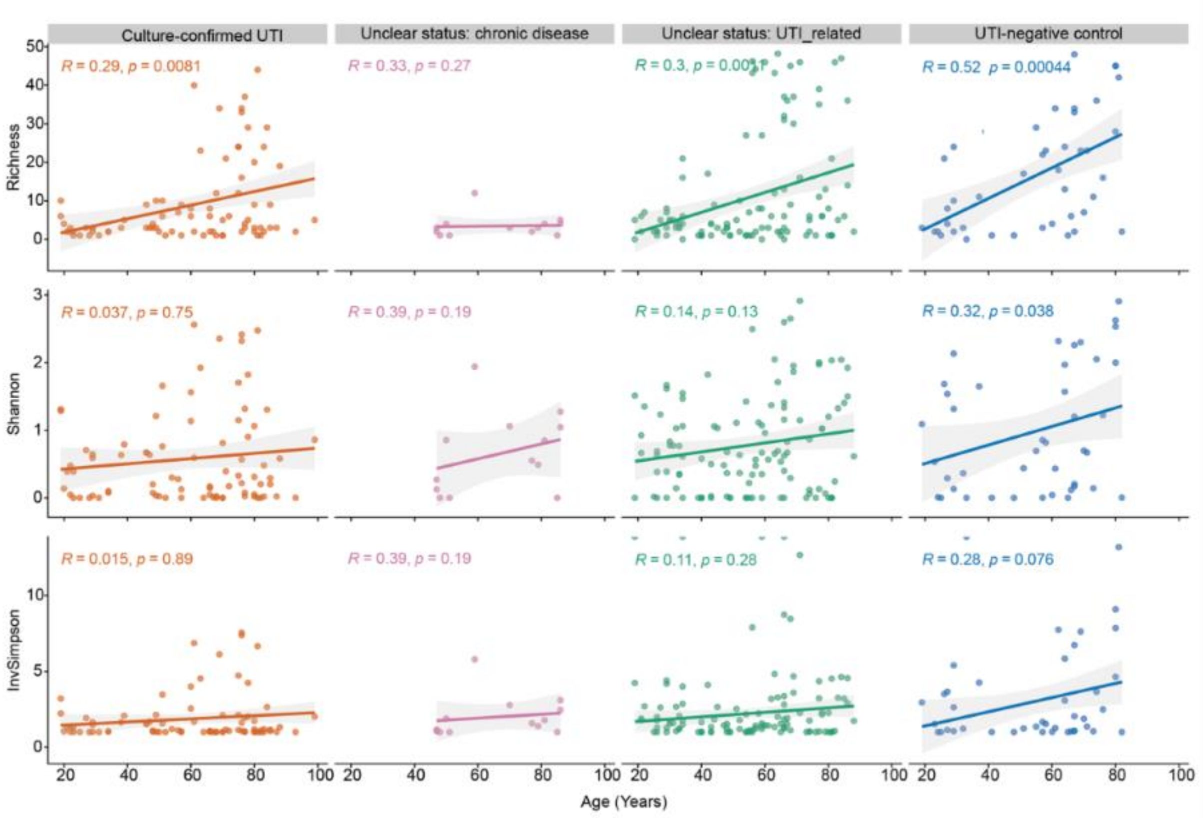


**Supplementary Figure 4. Richness and alpha diversity positive trends through age.** Distribution of the alpha diversity through age, divided by UTI status. The linear model was fitted to the alpha diversity of the samples and correlated with the age of every sample. Positive correlation was observed for the UTI negative group and the alpha diversity characterized by an increase of it through the age.

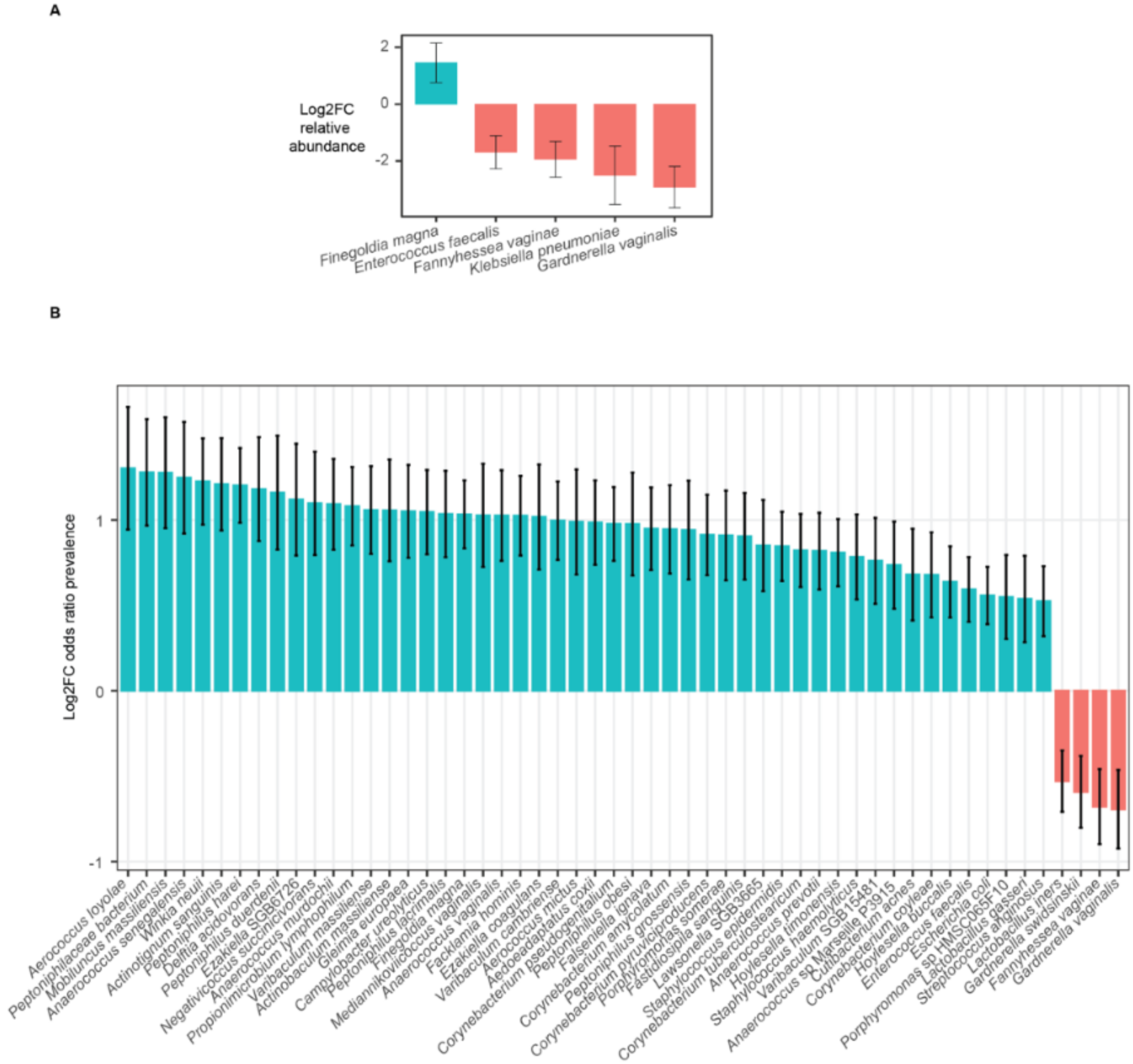


**Supplementary Figure 5. Differential abundance by age. A)** Differential abundance of bacterial species across age identified using MaAslin3. **B)** Differential prevalence of bacterial species across age, shown as log fold change (odds ratio).

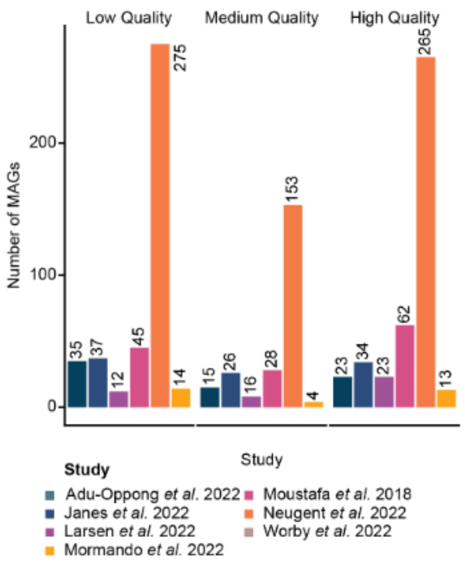


**Supplementary Figure 6**. **The number of low, medium, and high-quality MAGs were successfully recovered across all metagenomes.** High-quality genomes were recovered despite the low microbial biomass in many urine samples.

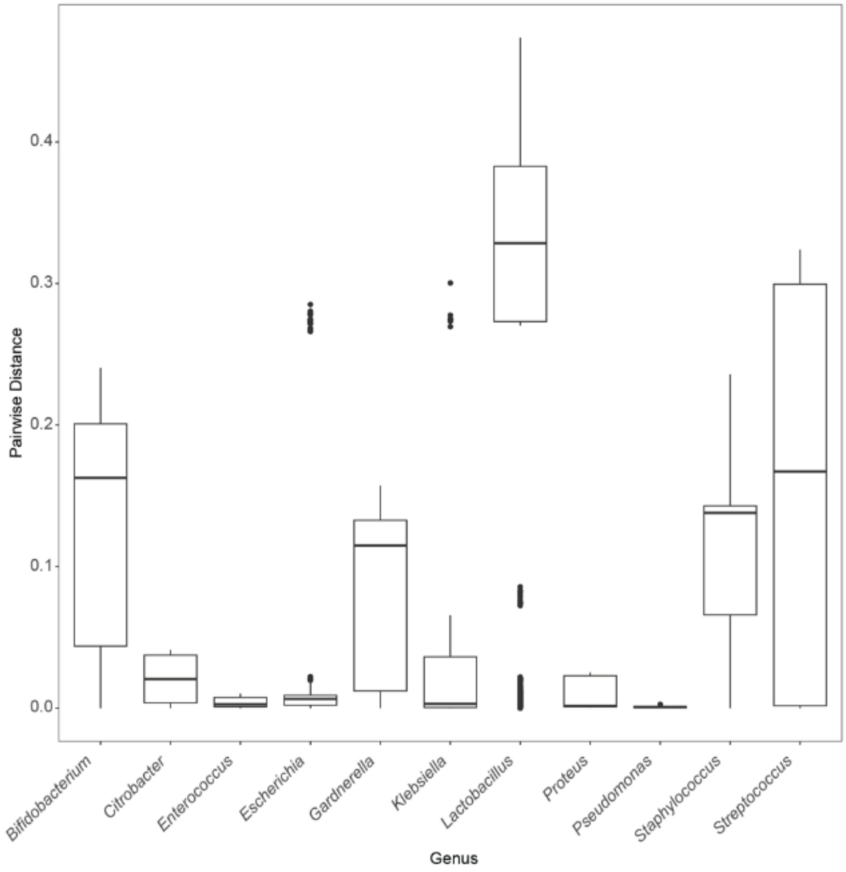


**Supplementary Figure 7. Bacteria commonly identified as pathogenic are clade-specific, while other taxa are phylogenetically diverse.** Distribution of phylogenetic distances within the main genera shown in **Fig. 4C**. Boxplots represent the range and variation of intra-genus phylogenetic distances, highlighting differences in lineage diversity across genera. Clinical relevance was determined based on clinical reports of most-prevalent bacteria causing UTI.

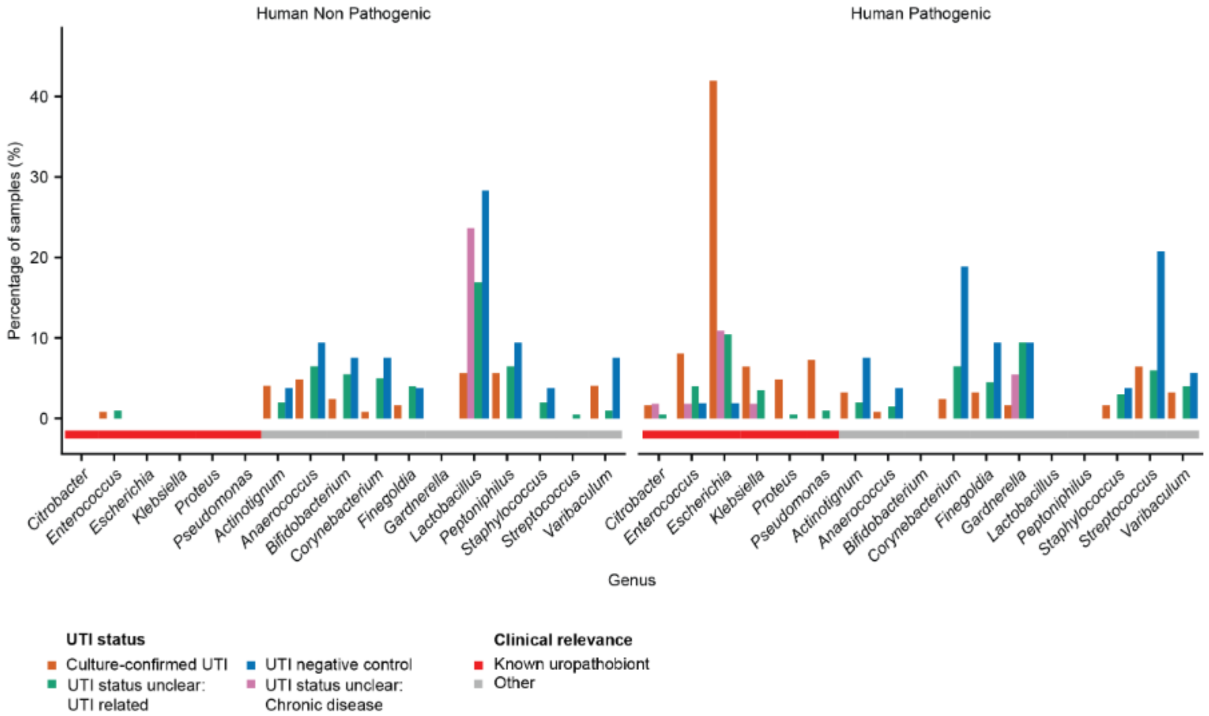


**Supplementary Figure 8. Prevalence of predicted pathogenic genera across UTI status groups**. Bar plot showing the prevalence of genera across all samples, grouped by predicted pathogenicity based on PathogenFinder2 [84] predictions. Genera are categorized as predicted human pathogens or non-pathogens. Non-common uropathobionts presented pathogenic potential in the negUTI group.

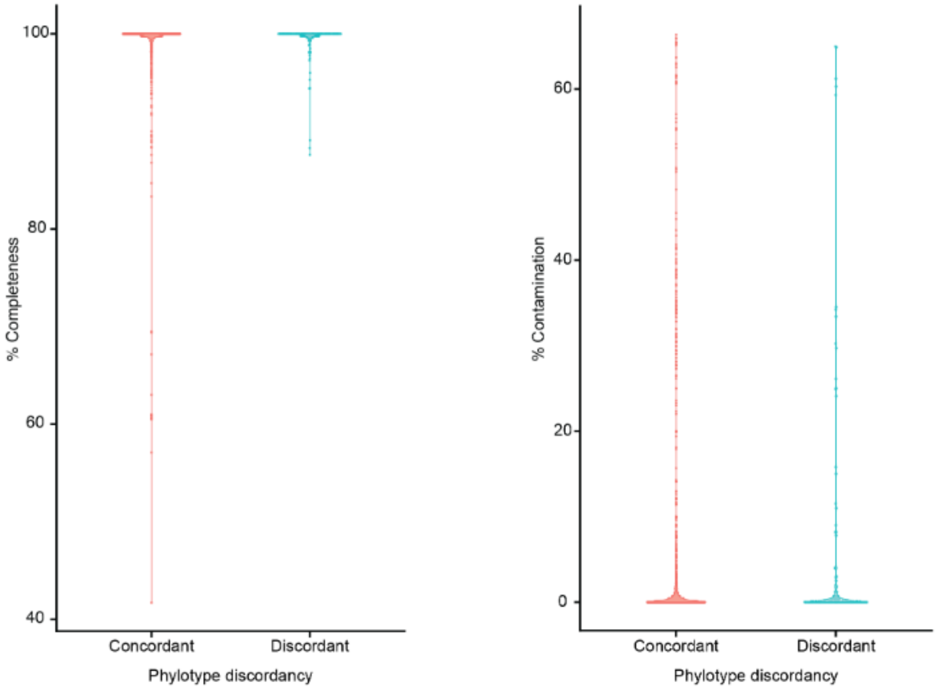


**Supplemental Figure 9. Genome quality does not explain phylotype discordances**. Violin plots showing completeness and contamination of *E. coli* MAGs grouped by concordance between ClermonTyping-assigned phylotypes and phylogenetic placement on the whole-genome tree. No significant differences were observed between concordant and discordant groups (Wilcoxon test, $p > 0.05$).

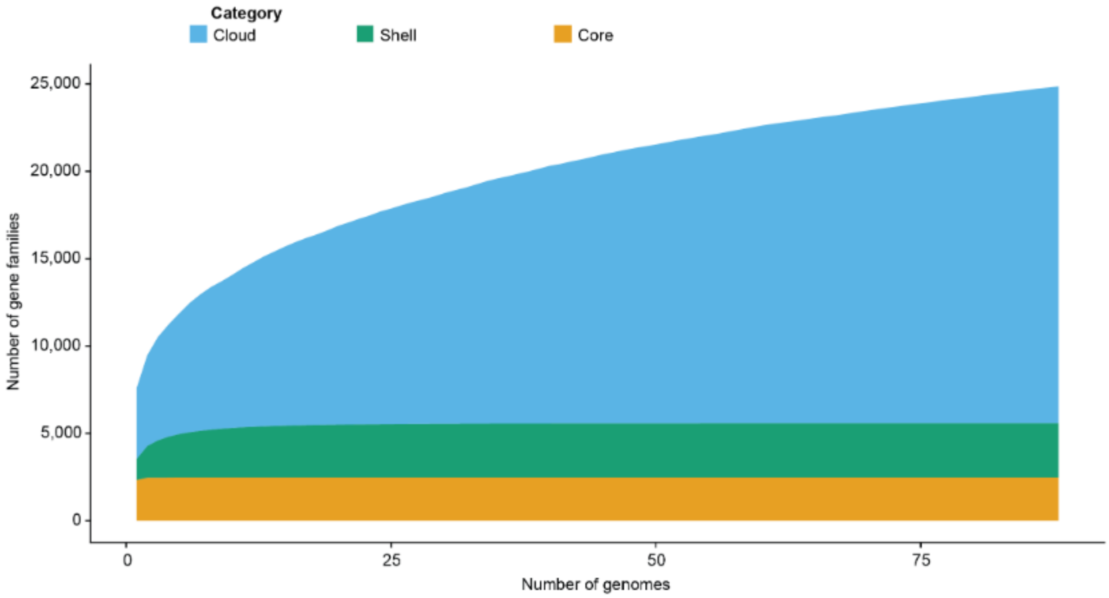


**Supplementary Figure 10. UTI-associated *E. coli* has an open pangenome.** Pangenome growth curve based on Heaps' law (α = 0.59) for only UTI positive *E. coli* genomes, indicating an open pangenome. Core and shell gene numbers stabilize, whereas cloud/accessory genes continue to increase with additional genomes being added.

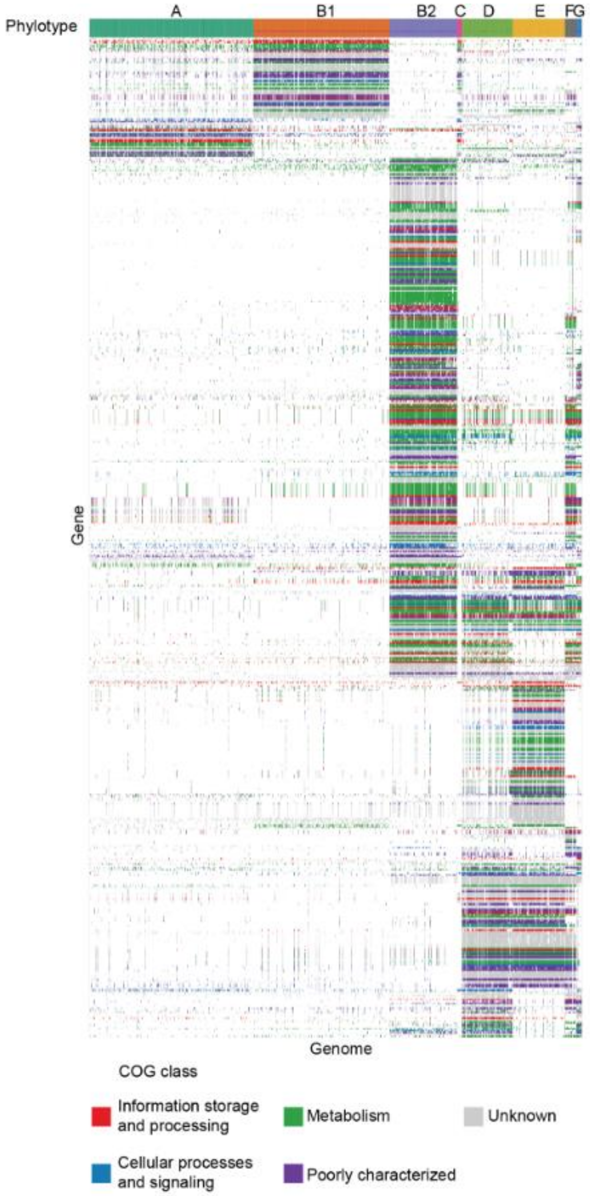


**Supplementary Figure 11. Genes detected by pan-GWAS to be significantly associated with phylotypes are related to various COG categories.** Heatmap of presence–absence for genes significantly associated with phylotypes identified in the pan-GWAS, colored by COG broad category.
